# Bmper is required for morphogenesis of the anterior and posterior semicircular canal ducts in the developing zebrafish inner ear

**DOI:** 10.1101/2021.06.27.450014

**Authors:** Sarah Baxendale, Esther C. Maier, Nikolaus D. Obholzer, Sarah Burbridge, Joseph Zinski, Francesca B. Tuazon, Nicholas J. van Hateren, M. Montserrat Garcia Romero, Mar Marzo, Kazutomo Yokoya, Robert D. Knight, Sean G. Megason, Mary C. Mullins, Tanya T. Whitfield

## Abstract

BMP signalling is known to have a conserved function in development of the semicircular canal system of the vertebrate inner ear, but its regulation, target genes and effects on cell behaviour during otic morphogenesis are not fully understood. We have characterised the effects of mutations in the zebrafish gene *bmper,* which codes for a regulator of BMP signalling with both pro- and anti-BMP roles in different developmental contexts. The inner ears of *bmper* mutant embryos develop with truncations of the anterior and posterior semicircular canal ducts. To image the developing ear in live embryos, we have exploited a new transgenic line, *Tg(smad6b:EGFP)*, which exhibits strong GFP expression in the otic epithelium. Morphometric analysis indicates defects in the *bmper* mutant ear from early stages of semicircular canal formation, correlating with a specific reduction in BMP signalling activity and specific loss of *dlx5a* expression in dorsal otic epithelium. Subsequent changes to cell shape occur at the truncation site and the dorsolateral septum. The *bmper* mutations that we describe are adult viable; truncation of the anterior and posterior semicircular canal ducts persists into adulthood. Our results argue against a major role for Bmper in specification of the pre-placodal region, induction of the otic placode, or development of the neural crest, processes in which Bmper function has previously been implicated. Instead, we conclude that a key requirement for Bmper function in the zebrafish is to promote BMP signalling during patterning and morphogenesis of the semicircular canal system.

## Introduction

Formation of the interlinked ducts and chambers that make up the semicircular canal system of the vertebrate inner ear is a spectacular example of epithelial morphogenesis in the developing embryo (reviewed in Alsina and Whitfield, 2017). BMP signalling is known to play an important role in otic patterning and morphogenesis at multiple stages of ear development, including in formation of the semicircular canal system. At gastrula stages, a dorsal (low)-to-ventral (high) gradient of BMP signalling activity patterns the dorsoventral axis of the entire embryo, a process that contributes to the positioning of the preplacodal region (PPR) and neural crest at the border between the neural plate and non-neural ectoderm (Nguyen et al., 1998; Tuazon and Mullins, 2015). The PPR gives rise to all cranial sensory placodes, including the otic placode, precursor of the inner ear. Correct PPR specification and placode induction involves modification of the BMP gradient; specifically, BMP signalling must be attenuated for otic placode induction within the PPR (Bhat et al., 2013; Reichert et al., 2013; reviewed in Groves and LaBonne, 2014).

After otic vesicle formation, genes coding for BMPs, BMP receptors and BMP signalling modulators show conserved patterns of expression in the developing ear, including in the semicircular canals and their associated sensory cristae (Chang et al., 2002; Chang et al., 2004; Morsli et al., 1998; Mowbray et al., 2001; Oh et al., 1996; Shawi and Serluca, 2008; Wu and Oh, 1996). In zebrafish, both *bmp2b* and *bmp4* are expressed at the anterior and posterior of the otic vesicle at 24 hours post fertilisation (hpf), and later in the developing cristae; *bmp7a* is expressed in the posterior of the otic vesicle, whereas *bmp7b* is expressed later in the outgrowing epithelial projections that fuse to form pillars of tissue (the hubs of the canal ducts) spanning the otic lumen, becoming down-regulated on fusion (Mowbray et al., 2001; Schmid et al., 2000; Shawi and Serluca, 2008). At least five genes coding for antagonists or modulators of BMP signalling (*bmper*, *fsta*, *chordin*, *smad6a*, and *smad6b*) are also expressed in or around the developing zebrafish ear (Esterberg and Fritz, 2009; Hammond and Whitfield, 2006; Kudoh et al., 2001; Mowbray et al., 2001). A transgenic reporter of BMP activity reveals expression in the otic vesicle, in a similar pattern to that of *bmp2b* at 24 hpf (Collery and Link, 2011).

As BMP signalling plays a central role in dorsoventral axial patterning of the embryo during early development, leading to embryonic lethal phenotypes for mutations in several BMP pathway genes, conditional approaches have been employed to study its function during inner ear morphogenesis. In the zebrafish, loss of *bmp2b* function during larval stages, following RNA-mediated rescue of embryonic axial patterning in *bmp2b* mutants, resulted in a complete absence of all three semicircular canal ducts in the adult ear, although ampullae, sensory cristae and the crus commune were present. These *bmp2b* mutant fish also displayed an abnormal swimming behaviour, indicative of balance defects (Hammond et al., 2009). In the mouse, conditional loss of *Bmp4* led to a loss of all three cristae and semicircular canal ducts in the most severe cases (Chang et al., 2008), and conditional inactivation of the *Alk3* type I BMP receptor gene in an *Alk6+/-* background resulted in thin or truncated canals (Ohyama et al., 2010). In chick, application or mis-expression of the BMP antagonists Noggin, DAN or Smad6 resulted in malformed or missing cristae and canals (Chang et al., 1999; Chang et al., 2008; Gerlach-Bank et al., 2004; Gerlach et al., 2000; Ohta et al., 2010). BMP signalling has been shown to drive the epithelial thinning that forms the dorsolateral pouch (precursor of the anterior and posterior semicircular canal ducts) (Ohta et al., 2010), and is thought to work via both canonical (SMAD-dependent) and non-canonical (PKA-dependent) pathways to regulate expression of *Dlx5* and *Hmx3*, respectively, in the chick otic vesicle (Ohta et al., 2016).

In this paper, we identify and characterise recessive mutations in the zebrafish *BMP-binding endothelial regulator* (*bmper*) gene, which codes for a BMP regulator and is itself transcriptionally responsive to BMP signalling. The zygotic *bmper* mutant displays very specific and fully penetrant but variably expressive defects in the inner ear, leading to truncations of the anterior and posterior semicircular canal ducts. We have imaged the otic phenotype using a new transgenic line, *Tg*(*smad6b:EGFP*), which provides a live readout for the expression of *smad6b*, a BMP inhibitor gene. Based on transcriptional changes and a reduction in BMP pathway activity in the *bmper* mutant ear, we propose a pro-BMP role for Bmper in dorsal otic epithelium, with *dlx5a* as a target gene. Both zygotic and most maternal-zygotic (MZ) *bmper* mutants are adult viable and fertile; we did not find any major defects in embryonic development of the neural crest or cardiovascular systems, tissues in which a role for Bmper has previously been suggested. Our work confirms the importance of BMP signalling in formation of the vertebrate semicircular canal system, and identifies Bmper as a key regulatory BMP-promoting factor in zebrafish dorsal otic morphogenesis.

## Materials and Methods

### Animals

Zebrafish lines used were AB, London wild type (LWT), King’s College wild type (KWT), *bmper^sa108^*, *bmper^del2^*, *bmper^del5^* (this work), *Tg(Xla.Eef1a1:h2b-mRFP1)* (Rodríguez-Aznar et al., 2013), *Tg(smad6b:EGFP-CAAX)* (this work) and *Tg(BMPRE:mRFP)* (Reichert et al., 2013). All mutants described are homozygous for the zygotic allele, except where maternal-zygotic (MZ) mutants were used, as indicated. Animals were maintained according to recommended husbandry regulations (Aleström et al., 2020), and all animal experiments conformed to UK Home Office or US regulations.

### Generation of the *Tg(smad6b:EGFP-CAAX)* line

The *Tg(smad6b:EGFP-CAAX)* line (referred to throughout as *Tg(smad6b:EGFP)*) was generated in an enhancer trap screen by ligating a 4.1 kb fragment upstream of the zebrafish *en2a* coding region to a minimal *c-fos* promoter driving membrane-(CAAX)-tagged EGFP in the Gateway vector pDestTol2. The construct was injected into the KWT strain using the Tol2 system (Kawakami, 2007). Injected embryos were grown up to adulthood and their offspring selected on the basis of GFP expression in the ear. Transgenic individuals were crossed to the *bmper^sa108^* line for imaging the otic epithelium.

### Identification of the *Tg(smad6b:EGFP)* line insertion site

Two methods were used to identify the insertion site of the transgene (Supplementary Figure S2). Thermal asymmetric interlaced (TAIL)-PCR (Liu and Whittier, 1995; Parinov et al., 2004), which alternates a higher annealing temperature for the specific Tol2 transgene primers, and a lower annealing temperature for degenerate primers to bind to the genomic sequence, was used to generate short fragments flanking the transgene insertion site. TAIL-PCR products from the secondary and tertiary nested PCR reactions were separated by gel electrophoresis, and fragments with the correct size shift—corresponding to the distance between the nested Tol2 primer sites—were isolated and sequenced. Tol2 and degenerate (AD) primer sequences used were: Tol2 5′-1, GGGAAAATAGAATGAAGTGATCTCC, Tol2 5′-2 GACTGTAAATAAAATTGTAAGGAG, Tol2 5′-3 CCCCAAAAATAATACTTAAGTACAG, Tol2 3′-1 CTCAAGTACAATTTTAATGGAGTAC, Tol2 3′-2 ACTCAAGTAAGATTCTAGCCAGA, Tol2 3′-3 CCTAAGTACTTGTACTTTCACTTG, AD-3 WGTGNAGNANCANAGA, AD-5 WCAGNTGWTNGTNCTG, AD-6 STTGNTASTNCTNTGC, AD-11 NCASGAWAGNCSWCAA.

Sequences from either side of the insertion site were mapped to the zebrafish GRCz11 genome assembly and two fragments (Tol2 5′-3/AD11 and Tol2 3′-3/AD5) identified the same locus at Ch18:19,713,380 (GRCz11) at the 3’ end of the *smad6b* locus. This insertion site was confirmed using the Targeted Locus Amplification (TLA) method (Cergentis B.V., Utrecht, Netherlands, (De Vree et al., 2014; Hottentot et al., 2017). For cell preparation, the protocol was adapted for use with zebrafish as follows (see Supplementary Fig. S2). Two hundred zebrafish embryos at the 15-20 somite stage were dissociated in a clean Petri dish under a coverslip in a minimal volume of calcium-free Ringer’s solution, as previously described (Baxendale et al., 2009). Cells were collected in a 1.5 ml reaction tube and separated from yolk platelets by centrifugation at 300g for 1 minute. The pellet of embryo cells was gently resuspended in Accumax cell dissociation medium and incubated until cells were fully dissociated. DNA crosslinking using ∼10^7^ cells and the Cergentis kit were used to fix the cells before being shipped to Cergentis B.V. for TLA analysis.

Two integration sites were found: one at Ch18:19,713,380 (GRCz11), corresponding to the *smad6b* locus, and a second insertion site at Ch16:50,170,719 (GRCz11) within the *cysteine-rich venom protein natrin-1-like* (*crvpn1l*) gene. Both insertion sites were confirmed by PCR using genome-specific primers to either side of the insertion: *smad6b* locus: F: GGGTTAGGGGTAGGAAAGGAATA, R: GCAAACATACCCACGTTGCTAT, *crvpn1l* locus: F: GGACATATCACCTAAATCCGCTG, R: GTACAATAAATACACCTCAATG.

### Mapping and identification of the genetic lesion using RNAseq and RNAmapper

To map and identify the *bmper* mutation, we followed the RNAmapper pipeline (http://www.rnamapper.org/) (Miller et al., 2013). In brief, mRNA was isolated from *bmper* mutant and sibling larvae at 3 days post fertilisation (dpf), sorted by ear phenotype into separate pools of 35 larvae each. Each pool was then sequenced to approximately 5 million PE76 reads on a HiSeq2000 machine, covering 61.7% of the transcriptome (Zv9.65) >4-fold. We obtained coverage for >95% of genes, with an average coverage of 26 reads per position. Sequencing reads were entered into the RNAmapper pipeline in FASTQ format. Based on de novo SNPs, RNAmapper predicted that the mutation lay within a 13.9 Mb interval on chromosome 16 (chr16: 1,444,165-15,420,254; peak AF: 0.951). Within the critical interval, we detected four candidate genes with potentially serious deleterious changes: *foxo6*, *stx12l*, *bmper* and *ptpn2a*.

Based on the known expression pattern of *bmper* in the otic vesicle and the known conserved role of BMP signalling in otic development, we prioritised *bmper* and confirmed a G>A transition mutation at 16:9261799 (Chromosome 16: 7,154,266-7,195,000 on GRCz11) by RT-PCR and sequencing.

### TALEN mutagenesis

TALEN targeted mutagenesis was used to create a 2 bp and 5 bp deletion at 669/2004 (223/668aa) in exon 8 of the 16-exon transcript (ENSDARG00000040893) (Fig. 3).

### In situ hybridisation and Immunohistochemistry

in situ hybridisation was performed as previously described (Oxtoby and Jowett, 1993; Thisse and Thisse, 2008). Probes used are listed in Supplementary Table S1. Whole embryos were fixed in 4% paraformaldehyde overnight and stained with anti-phospho-Smad1/5/9 Rabbit monoclonal antibody (Cell Signaling Technology #13820S; 1:500), anti-phospho-H3 Rabbit monoclonal antibody (Cell Signaling Technology #9701S; 1:500), followed by Goat anti-Rabbit Alexa 568 secondary antibody (1:200).

### Alcian blue staining

Craniofacial cartilages were stained using the alcian blue method, as previously described for zebrafish (Felber et al., 2015).

### Imaging

Fixed embryos stained by in situ hybridisation or with alcian blue were imaged on an Olympus BX51 compound microscope equipped with DIC optics, a Camedia C-3030ZOOM camera and CELL-B software. For Figures S4, 5 and 6, batches of embryos from a mixed cross of homozygous and heterozygous *bmper* mutant fish, with an expected ratio of 50:50, homozygous mutant:heterozygous sibling, were used. >30 embryos were analysed with each probe and at each stage of development. Fluorescent antibody stains were imaged on an Olympus FV-1000 confocal microscope. Live transgenic embryos were anaesthetised in 0.5 mM MS222 (3-aminobenzoic acid ethyl ester) and mounted in 3% methylcellulose or in 0.7% agarose for imaging with a 20× objective on an Olympus FV-1000 confocal microscope or a Zeiss Z.1 light-sheet microscope. Light-sheet datasets were analysed in Zen black (Zeiss) or arivis software. Fiji (Schindelin et al., 2012) or Adobe Photoshop were used to adjust brightness and contrast. Levels of P Smad5 staining in Fig. 5 were quantified as previously described (Zinski et al., 2019).

### Adult ear dissections and TRITC-phalloidin staining

Adult zebrafish were killed with an overdose of MS222 (3-aminobenzoic acid ethyl ester) and decapitated. Heads were fixed in 4% paraformaldehyde for at least 1 week. Ears were dissected and stained with TRITC-phalloidin (Sigma) as previously described (Baxendale and Whitfield, 2016). The image shown in Fig. 7G appeared in this publication.

### Data Analysis

Statistical analyses were performed using GraphPad Prism version 9.

## Results

### The zebrafish *bmper* mutant ear has truncated anterior and posterior semicircular canal ducts

We identified a recessive mutation affecting ear morphology in the background of the zebrafish *otx1b^sa96^* line, obtained from the Sanger Centre Zebrafish Mutation Project (Kettleborough et al., 2013). The mutation, named *bmper* following the identification of the genetic lesion (see below), results in a truncation of the anterior and posterior semicircular canal ducts, with defects visible under the dissecting microscope from around 52 hours post fertilisation (hpf). Other structures in the ear, including the otoliths, all three semicircular canal cristae and the lateral (horizontal) semicircular canal duct, appear to develop normally (Fig. 1A, Supplementary Movies 1-3). The epithelial projections that meet and fuse to form the pillars around which the anterior and posterior semicircular canals form are all present, and fusion occurs at the correct time. However, the pillars are located closer to the dorsal otic epithelium within the otic vesicle and the dorsolateral septum (DLS) that separates the anterior and posterior canals does not form correctly (Fig. 1B). The mutation does not appear to affect development of other organs or the size or length of the embryo significantly (Fig. 1C), and homozygous mutants are adult viable. The ear phenotype appears to be almost fully penetrant (23.8% of embryos showed the ear phenotype from a cross between heterozygous parents; *n*=734), but there is some variability in expressivity: in a minority of mutant embryos, one or both ears had a weaker phenotype, with one canal, usually the posterior, unaffected and of normal size. In a few embryos, the truncations were more severe (Fig. 1D, Supplementary Fig. S1).

**Figure 1.**
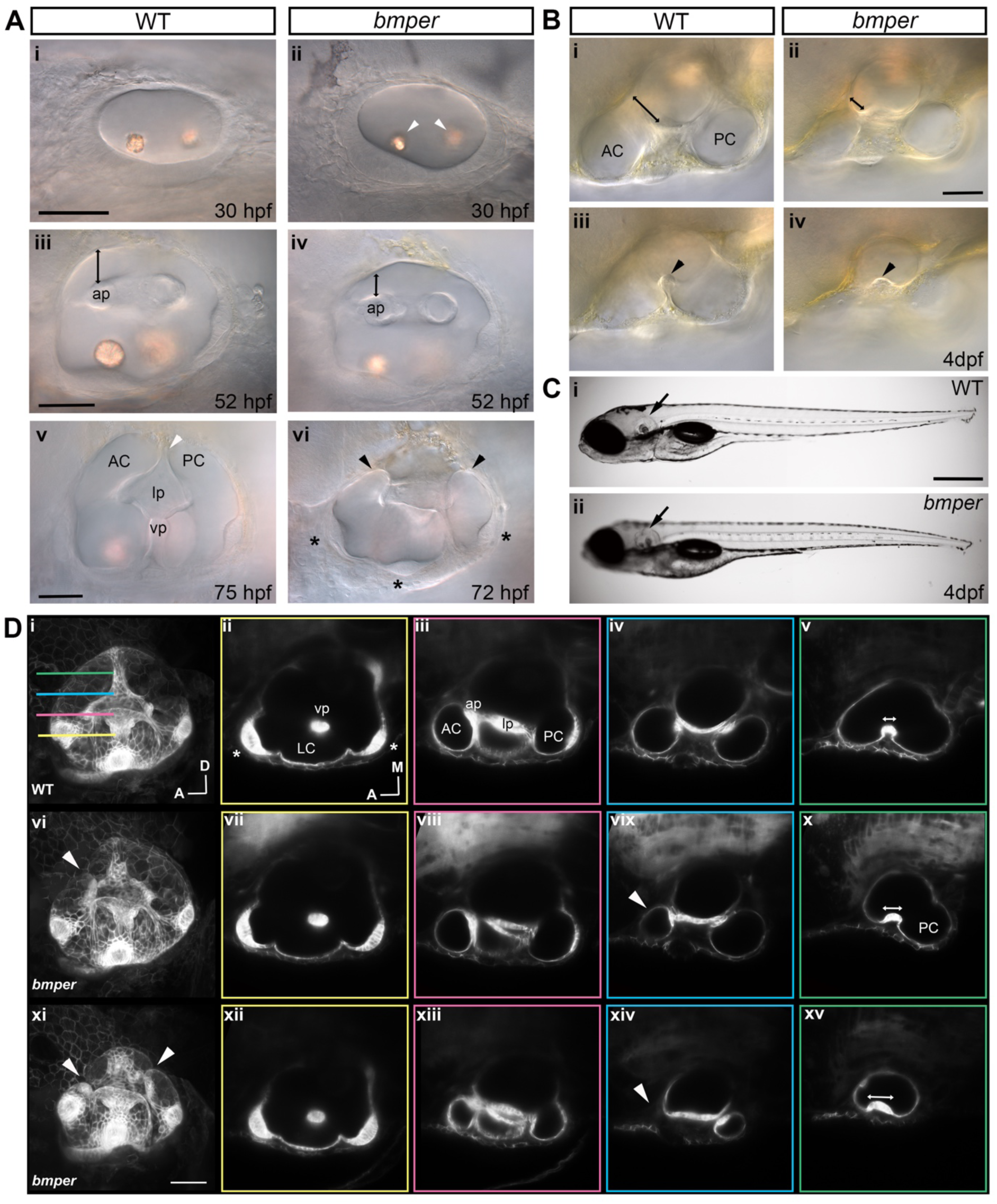
The ears of *bmper* mutant embryos have truncated anterior and posterior semicircular canal ducts. **A** (i-vi) Live DIC images of the otic vesicle in phenotypically wild-type (WT) sibling (i, iii, v) and *bmper^sa108^* (ii, iv, vi) embryos, lateral view, anterior to the left. (i, ii) 30 hpf, no clear differences in otic vesicle size or shape are seen, and otoliths form normally in *bmper* mutants (white arrowheads). (iii, iv) 52 hpf, *bmper* mutants have a smaller gap between the anterior and posterior epithelial pillars that span the vesicle and the dorsal edge of the vesicle (black lines above anterior pillar). (v, vi) 75 hpf (sibling) and 72 hpf (*bmper*); the semicircular canal lumens have formed in wild-type sibling embryos (v), and the anterior (AC) and posterior (PC) canal ducts span the full height of the vesicle, separated by the dorsolateral septum (DLS, white arrowhead). Note the ventral pillar (vp) can be seen below the lateral projection (lp). In *bmper* mutants, the anterior and posterior canal ducts are truncated below the height of the otic vesicle (black arrowheads); the ventral pillar is present but cannot be seen in this view. All three cristae are present (black asterisks). **B** (i-iv) Dorsal view of the otic vesicle, anterior to the left, dorsal to the top, at 4 dpf in wild-type sibling (i, iii) and *bmper* mutant (ii, iv). The wild-type sibling shows clear anterior (AC) and posterior (PC) canal lumens (i) and a pronounced DLS (black arrowhead, iii), whereas in *bmper* mutants the anterior canal lumen has a smaller diameter and the length of the anterior pillar (black line) is shorter than in the wild type (ii); the DLS is also shorter and wider (black arrowhead). **C** (i, ii) Live DIC image of 4 dpf wild-type sibling (i) and *bmper* mutant (ii) larvae. Arrows mark the otic vesicle (mis-shapen in *bmper* mutants). Body length, pigmentation pattern and swim bladder inflation appear normal in *bmper* mutants. There was no significant difference in body length at 3.5 dpf (two-tailed *t*-test, *p*=0.427, *n*=29) **D** (i-xv) Light-sheet microscope images using the *Tg(smad6b:GFP)* line, showing a Maximum Intensity Projection (MIP) lateral view of 4 dpf otic vesicles on the left (i, vi, xi) and corresponding single *z* slices of a dorsal view through four different planes for each otic vesicle. (i-v) WT, wild-type sibling, (vi-xv) *bmper* mutant. Each *z* slice is taken at the level shown by the coloured lines on the WT lateral view (i), such that the *z*-slices in the yellow boxes (ii, vii, xii), are taken at the lowest ventral plane and the *z*-slices in the green boxes are taken at the most dorsal plane. The distance between each of the four slices is 26.8 μm (*z* interval=1.381 μm; sample images are 20 *z* planes apart). vi, xi, lateral views of two different *bmper* mutants: (vi) has a weaker phenotype with a truncation in the anterior canal (white arrowhead) and a complete posterior canal; (xi) has a more severe phenotype with both anterior and posterior truncated canals (white arrowheads). The dorsal views show the cristae (white asterisks), ventral pillar and lateral canal form normally in both *bmper* otic vesicles shown (yellow planes); note the more severe *bmper* otic vesicle is slightly smaller (xi). The next plane (magenta) is at the level of the lateral projection and the anterior pillar in all four otic vesicles; the lumen of the anterior canal is smaller in both *bmper* mutants (viii, xiii). The two more dorsal planes (blue, green) clearly show the truncated anterior canal in both *bmper* mutant ears (white arrowhead, vix, xiv) compared to the wild type (iv). In the weaker *bmper* phenotype the posterior canal remains intact (x), but the DLS does not develop normally in either *bmper* mutant (white double-headed arrow). Abbreviations: AC, lumen of anterior semicircular canal; ap, anterior pillar; PC, lumen of posterior semicircular canal; pp, posterior pillar; vp, ventral pillar; lp, lateral projection; DLS, dorsal lateral septum; A, anterior; D, dorsal; M, medial, MIP, maximum intensity projection. Scale bars: 50 μm in Ai, for Aii, Aiii, for Aiv, Av, for Avi; 50 μm in Bii, for Bi; 500 μm in Ci, for Cii, iii, iv; 50 μm in Dxvi for Di-Dxx.

To image the semicircular canal duct abnormalities in the zebrafish ear, we crossed heterozygote carriers of the mutation to a transgenic line expressing GFP in the otic epithelium. This line was generated by random insertion of a construct consisting of a fragment of *en2a* upstream sequence and a minimal *c-fos* promoter driving expression of membrane-tagged EGFP (Supplementary Fig. S2); serendipitously, some transgenic fish showed GFP expression in the ear, and were isolated on this basis. Using a TAIL PCR-based approach (Liu and Whittier, 1995) and Targeted Locus Amplification (TLA, Cergentis), we identified two separate genomic sites of insertion of the transgene: one in the 3’ region of the *smad6b* locus on Chromosome 18, and a second site on Chromosome 16 in the *cysteine-rich venom protein natrin-1-like* (*cvrvpn1l*) locus (see Materials and Methods and Supplementary Fig. S2). The GFP expression pattern in the transgenic line, which we have named *Tg(smad6b:EGFP*), recapitulates the endogenous expression pattern of the inhibitory SMAD gene *smad6b*, including strong expression in the ear; endogenous *smad6b* expression is unaffected (Supplementary Fig. S2C). We do not see any GFP expression that correlates to the *crvpn1l* expression pattern as assayed by in situ hybridisation (Supplementary Fig. S2).

The *Tg(smad6b:EGFP)* embryos provide an excellent tool for imaging otic epithelia in the live zebrafish embryo from 24 hpf through to adulthood (Fig. 1D, Supplementary Fig. S1 and Supplementary Movie 2). Analysis of wild-type and *bmper* mutant embryos in the *Tg(smad6b:EGFP)* line at 96 hours post fertilisation (hpf) using light-sheet microscopy confirmed that the anterior and posterior canal ducts were truncated, forming a blind-ended stump, with the anterior canal often more strongly affected. Dorsal view optical sections showed that the three sensory cristae and ventral pillar developed normally, although the ears of embryos with strong *bmper* phenotypes were slightly smaller at this stage (Fig. 1D, Supplementary Movie 2). The space connecting the anterior and posterior canals, which will later become the crus commune, was still present and of a normal height in all *bmper* mutant ears; however, the dorsolateral septum (DLS), which bisects this area at the dorsal boundary between the anterior and posterior canals, was wider and flatter in *bmper* mutants (Fig. 1C, D).

### Live imaging reveals morphological and cellular changes in anterior and posterior canal formation

Imaging of the otic vesicle revealed changes in the overall shape at 55 hpf, after the anterior and posterior epithelial projections have grown in to the otic lumen and fused to form pillars (Fig. 1). We used 3D-rendered images of *bmper* mutant ears in the *Tg(smad6b:EGFP)* line to quantify any morphometric changes in 3D. The variability in the presence and size of the canal truncations made it challenging to characterise timing of onset of differences in ear shape. To orientate the ear so that all measurements were comparable, key features that were not affected in the mutant were used, including the endolymphatic duct (ED) and the three cristae.

Measurements were taken in lateral views of otic vesicle height (ED to ventral crista), width (anterior to posterior cristae), distance from ED to base of the anterior projection, distance between the anterior pillar and the dorsal otic epithelium, and distance across the base of the lateral projection. Changes were seen in *bmper* mutants with all measurements apart from height (Fig. 2). The *bmper* ear was slightly wider and the height above the anterior pillar was significantly reduced, as could be seen in the DIC images where shape of the top of the ear, normally domed, is more pointed in the *bmper* mutant (Fig. 1). Distance from the ED to the anterior projection was also reduced (either due to reduced height of the *bmper* otic vesicle, or to abnormal positioning of the anterior projection). One change that was only apparent from the fluorescent images is the width of the lateral projection. In wild-type embryos, the lateral projection grows into the otic lumen as a single evagination, bifurcating to form the anterior and posterior bulges that meet and fuse with the anterior and posterior projections, respectively. In *bmper* mutants, the lateral projection grows into the lumen, but the base of the projection is wider, and the anterior and posterior bulges are more widely separated. This combination of the wider lateral projection and the reduced gap above the anterior pillar means that the hub of the canal forms closer to the edge of the otic vesicle, and is a possible cause of the canal truncations.

**Figure 2.**
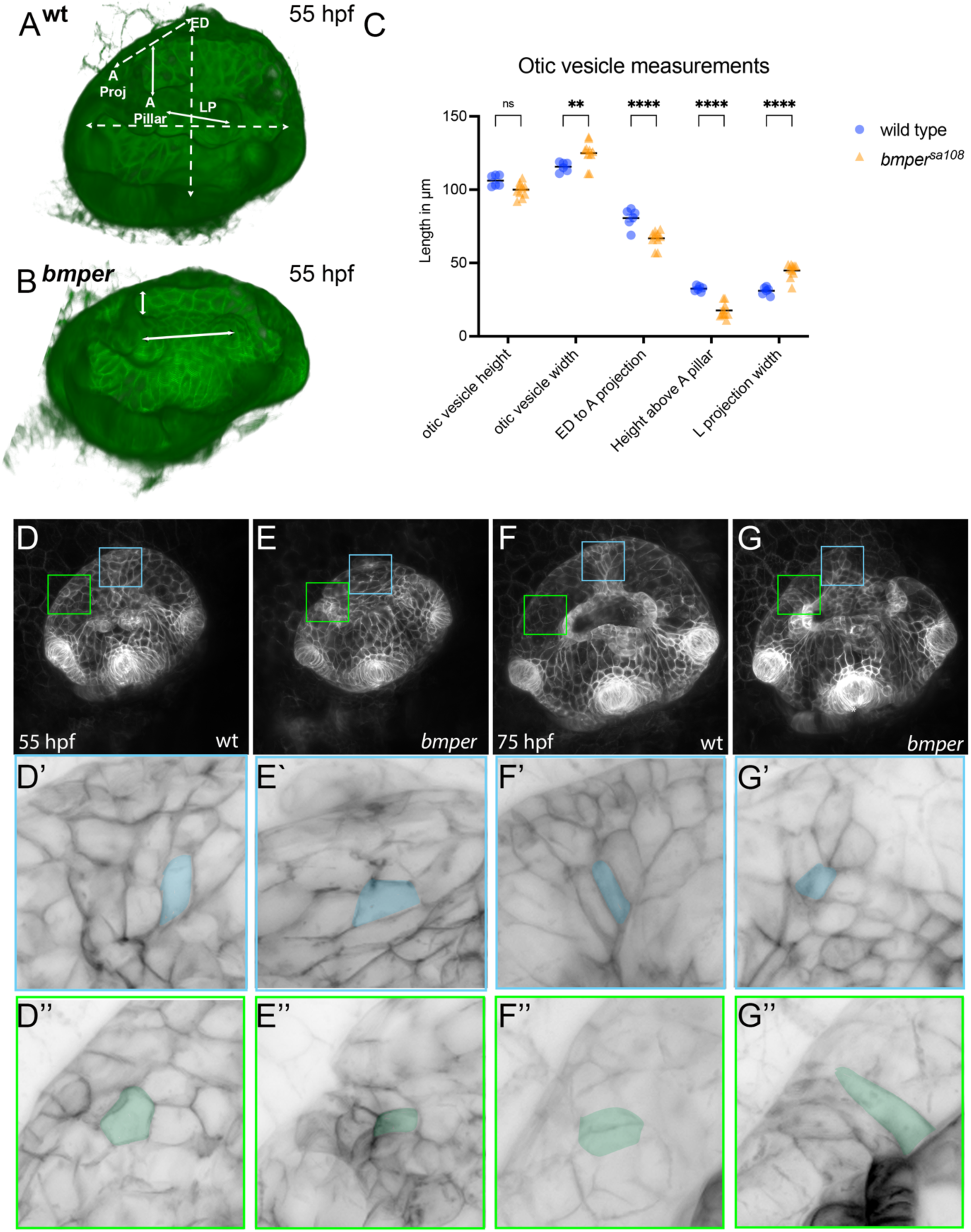
Live fluorescent imaging reveals changes in morphometric measurements and cell shape changes in *bmper^sa108^* embryos. **A–C.** Measurements of otic vesicle shape and features in 3D using transparency-rendered images of wild-type (A) and *bmper* mutant ear (B). *Tg(smad6b:EGFP*) embryos at 55 hpf taken with light-sheet microscopy. The internal otic vesicle structure is shown viewed from the medial edge of the ear looking towards the lateral edge, anterior to the left. Positions of the measurements taken are shown on the wild-type ear (A). White dashed lines correspond to otic vesicle height, width and the distince from the base of the ED to the base of the anterior projection. Continuous lines correspond to distance between the anterior pillar fusion site and the dorsal otic epithelium, and the width of the lateral projection (measured at the base of the lateral projection on the lateral side of the ear). **C.** Graph of the measurements in multiple ears for wild type (*n*=6 ears, *N*=3 fish) and *bmper* mutants (*n*=10, *N*=5). Each ear was imaged twice, from a dorsal and a lateral view, and each point on the graph represents the average of the two measurements. There were significant differences in all measurements apart from otic vesicle height. Two-way ANOVA with Šidák’s correction for multiple comparisons: \*\**p*=0.0066; \*\*\*\**p*<0.0001; ns, not significant. Abbreviations ED, endolymphatic duct; A, anterior; LP, lateral projection. **D–G’’**. Cell shape changes at the DLS and truncation points. Maximum Intensity Projections (MIPs) of *Tg(smad6b:EGFP)* in wild-type (D,F) and *bmper* mutant (E,G) ears at 55hpf (D,E) and 75 hpf (F,G). Blue boxes in D’–G’ correspond to the detailed view below of the DLS; green boxes in D’’–G’’ correspond to the detailed view of the site of the truncation. Manual segmentation of selected cells, pseudocoloured blue and green, highlighting the difference in cell shape between wild type and *bmper*.

The dorsolateral septum (DLS) dividing the anterior and posterior canal ducts at the dorsal part of the ear is also abnormal in *bmper* mutants. Analysis of the region dorsal to the lateral projection revealed changes in cell shape and organisation (Fig. 2). In wild-type embryos at 55 hpf, the cells above the lateral projection begin to elongate in the dorsoventral axis, and align to form the DLS. In *bmper* mutants, cells fail to align in an organised way. Cell shape changes were also found at the sites of the canal truncations. Here, cells appeared more tightly packed and were either smaller and rounded or elongated toward the truncation point in *bmper* mutants, whereas in the wild-type, cells on the outer wall of the canal have a squamous morphology (Fig. 2). We also noted that in *bmper* mutants with a weaker phenotype, where the canal duct had not truncated, a band of elongated cells was still present at a region that is the likely site where a truncation would have occurred. This suggests that changes in cell shape observed in *bmper* mutants do not always result in the truncation of the canal.

### Identification of mutations that disrupt the *bmper* gene

To identify the genetic lesion underlying the mutant phenotype, we used RNAmapper (Miller et al., 2013) to search for candidate causative SNPs in the mutant exome. We found that the mutation maps to chromosome 16 roughly at position 10,000,000, or between 1,500,000 and 15,000,000 (Fig. 3A, Supplementary Fig. S3). Within this interval, we detected the gain of a STOP codon with 8-fold coverage at position 9261799 (tGg>tAg, or W>*) in the *BMP-binding endothelial regulator* (*bmper*) gene (ENSDARG00000101980 – GRCz11), formerly known as *crossveinless 2* (*cv2, cvl2*; *bmper* is the orthologue of *Drosophila crossveinless 2* (*cv-2*)). Sequencing of genomic DNA prepared from individual embryos (*n*=20) from a cross between heterozygous parents confirmed linkage of the G>A SNP with the ear phenotype (Fig. 3B). The Sanger Centre has reported the identical mutation in their collection (*bmper^sa108^*), which was derived from the same round of mutagenesis (E. Busch-Nentwich, personal communication), lending further support to our sequencing results. Presumably, the *bmper^sa108^* and *otx1b^sa96^* lines are descendants of the same mutagenised male; we therefore adopted the designation *bmper^sa108^* for the mutation.

**Figure 3.**
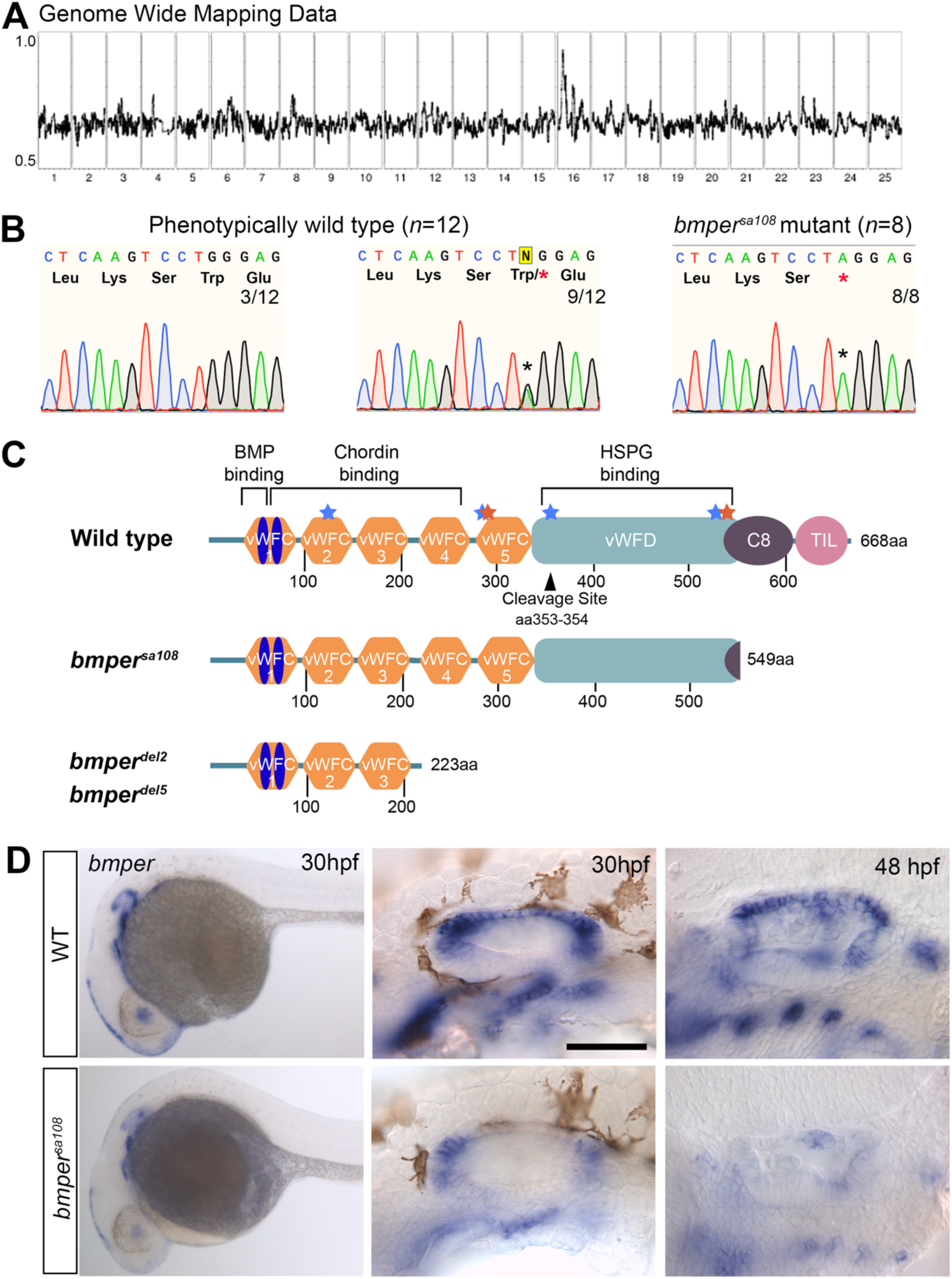
Position of mutations at the *bmper* locus and expression of *bmper* in wild-type and *bmper^sa108^* embryos. **A.** RNAseq genome-wide map data showing a strong peak on Chromosome 16. Numbers along the *x* axis indicate chromosomes, 1-25; the *y* axis shows the mutant marker allele frequency. **B.** Sequence traces from wild-type, heterozygous and homozygous *bmper* mutant embryos, taken from a *bmper* heterozygous cross. A G>A SNP changes a Tryptophan residue to a STOP codon in *bmper^sa108^* mutant embryos. **C.** Schematic of the zebrafish Bmper protein structure in wild-type and *bmper* mutant alleles. The protein contains five cysteine-rich von Willebrand factor type C domains (vWFC); a von Willebrand factor type D domain (vWFD); a cysteine-rich domain (C8), and a trypsin inhibitor-like domain (TIL). vWFC1 has two sub-domains shown in blue; the N terminal domain is involved in BMP binding and C terminal domain is part of the chordin-binding domain. The vWFD domain is suggested to bind to heparan sulphate proteoglycans (HSPG). Positions corresponding to human disease mutations associated with DSD are shown in blue (V137D, Q309*, P370L, C546*); mutations associated with the milder condition ISD are shown in orange (W314*, R558*). Positions are at the homologous position in the zebrafish protein sequence. Truncations associated with different alleles are shown. **D.** Expression of *bmper* in the inner ear of wild-type (top row) and *bmper^sa108^* mutant embryos (bottom row) at 30 hpf, 32 hpf, 48 hpf, anterior to the left. Strong expression can be seen in the dorsal otic vesicle that is down-regulated. Expression in *bmper^sa108^* mutant embryos is down-regulated throughout the embryo (left hand panels); right hand panels show enlargement of the otic region. Scale bar: 50 μm in view of detailed otic vesicle.

The Bmper protein sequence predicts a secreted glycoprotein containing five conserved cysteine-rich von Willebrand factor (vWF) type C domains. The STOP codon in the *bmper^sa108^* allele falls at amino acid position 549, within the vWFD domain. Interestingly, this is very close to the position of disease-causing mutations in the human gene (Fig. 3C)(Kuchinskaya et al., 2016). The mutation thus predicts a truncated protein in which the vWFC domains remain intact, but the uncharacterised cysteine-rich and conserved Trypsin Inhibitor domains towards the C terminus are lost (Fig. 3C). Importantly, the STOP codon is downstream of the predicted cleavage site required for conversion of Bmper from an anti- to a pro-BMP factor (Rentzsch et al., 2006), suggesting that the predicted protein product could retain some pro-BMP activity. We therefore compared *bmper^sa108^* to two further alleles, *bmper^del2^* and *bmper^del5^*, generated in one of our laboratories (MCM) by TALEN mutagenesis. These alleles have 2- and 5- base pair deletions, respectively; both result in a frame-shift leading to a STOP codon at amino acid position 223, between the third and fourth vWFC domains (Fig. 3C). Although the STOP codon in these *bmper^del2^* and *bmper^del5^* alleles is upstream of the predicted cleavage site, the ear phenotype is indistinguishable from that in *bmper^sa108^* mutants, and the mutations, like *bmper^sa108^*, are also adult viable (Supplementary Fig. S3).

### Expression of *bmper* mRNA in the developing zebrafish ear

Previous studies have described the expression pattern of the *bmpe*r transcript in the wild-type zebrafish embryo (Esterberg and Fritz, 2009; Kudoh et al., 2001; Kwon and Riley, 2009; Moser et al., 2007; Rentzsch et al., 2006). The otic vesicle is a major site of expression, which has not previously been studied in detail (Fig. 3D, Supplementary Fig S3). In wild-type embryos, expression is present in the posterior otic epithelium at the 12-somite stage (12S), later spreading anteriorly across the dorsal otic epithelium. At 24-30 hpf, expression is present throughout dorsal region, with higher expression in the anterior and posterior poles at 24 hpf, while *bmper* expression is absent in ventral otic epithelium. During outgrowth of the epithelial projections that form the semicircular canal system, *bmper* is expressed in the epithelium that forms the outer walls of the canal ducts, and at lower levels in the projections and bulges that fuse to form the pillars (the ‘hubs’ of each semicircular canal). By 2 dpf, expression appears more ventrally at the margins of the sensory cristae, and by 3 dpf, weak expression persists throughout the semicircular canal system, together with the support cell layer of the sensory maculae and cristae (Supplementary Fig. S3).

In *bmper^sa108^* mutants at 3 dpf, RNAmapper analysis detected a small (less than two-fold) but significant (*p*=0.002) reduction in expression of one of two *bmper* isoforms. As RNA for this analysis was taken at a stage when the semicircular canal duct truncation was already apparent, we verified this reduction in *bmper* mRNA levels in *bmper^sa108^* mutants at earlier developmental stages by in situ hybridisation (Fig. 3D). The observed systemic reduction in *bmper* transcript levels could be due to nonsense-mediated mRNA decay of the mutant transcript, or could reflect a reduction in BMP signalling activity, as *bmper* is a known transcriptional target of BMP signalling (Reichert et al., 2013; Rentzsch et al., 2006).

In the mouse, Bmper has been proposed as a candidate target gene for Tbx1, although Tbx1 binding sites in the murine *Bmper* gene were not reported to be conserved with the zebrafish (Castellanos et al., 2014). We examined the expression of *bmper* in *tbx1^-/-^* mutant embryos, which have a severe ear phenotype, with absent or vestigial epithelial projections (Radosevic et al., 2011; Whitfield et al., 1996), but did not see major differences in the levels of *bmper* expression (Supplementary Fig. S3E). Expression of *bmper* that normally marks the cristae in the ventral part of the ear was lost, but this is to be expected, as cristae do not develop in the *tbx1^-/-^* mutant ear. *bmper* expression levels were also unaffected in *sox10^-/-^* mutant embryos (Supplementary Fig. S3).

### Expression of otic markers in *bmper^sa108^* mutants reveals specific defects in dorsal otic epithelium

To understand the basis of the otic phenotype in *bmper* mutants further, we examined expression of other otic markers in the ear of *bmper^sa108^* mutant embryos over a range of developmental stages using in situ hybridisation. We first focussed on orthologues of *Dlx5* and *Hmx3*, known to be targets of BMP signalling in the developing chick ear (Ohta et al., 2016). In zebrafish, *dlx5a* and *hmx3a* expression domains overlap with that of *bmper* in the dorsal otic vesicle from 24 hpf to 48 hpf (Fig. 4). We found robust changes in the expression pattern of *dlx5a*, which was prematurely lost in dorsal otic epithelium from 30-48 hpf, initially at the anterior and posterior poles, but retained in the cristae and ED (Fig. 4). Expression of *dlx5a* at earlier (27 hpf) and later (72 hpf) stages did not show any changes in level or spatial extent in *bmper^sa108^* mutant embryos (Fig. 4). The wild-type otic expression pattern of zebrafish *hmx3a* is complex at 40 hpf, with discrete bands of expression in the anterodorsal and posterodorsal domains, but lacking in the dorsal region where the ED will form and a in small anterior gap (Fig. 4) (Hartwell et al., 2019). Quantification of the spatial extent of *hmx3a* domains of expression revealed subtle and transient differences in the *bmper* mutant ear, although levels appeared unaltered. The anterodorsal band of *hmx3a* expression was reduced in *bmper^sa108^* mutants, and the dorsal gap in expression, which includes part of the same region that lacks *dlx5a* expression, was expanded (Fig 4). By 72 hpf, however, the *hmx3a* otic expression pattern in *bmper^sa108^* mutant embryos was indistinguishable from the wild type (Fig. 4).

**Figure 4.**
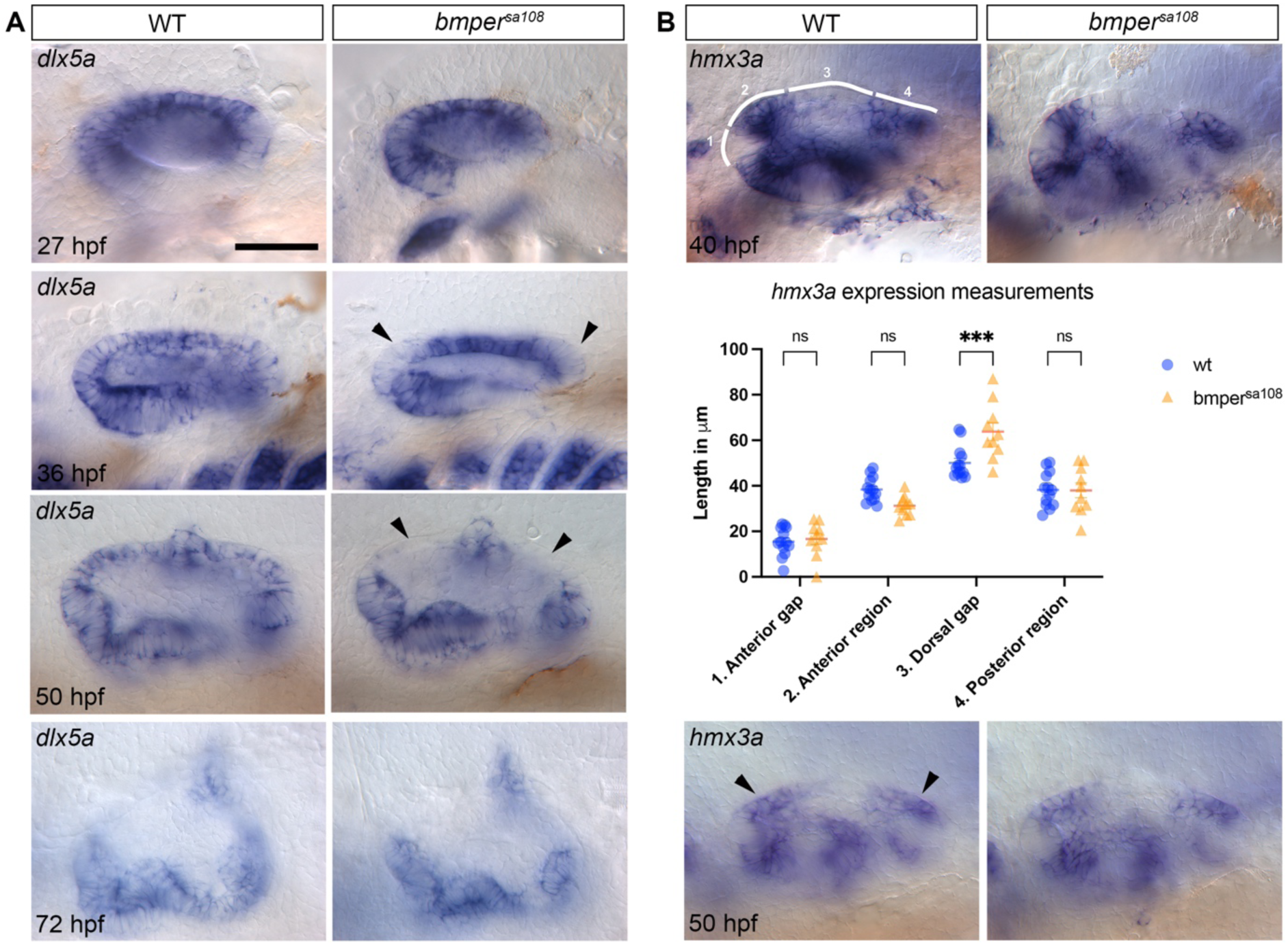
*bmper^sa108^* mutants have a specific loss of *dlx5a* in the dorsal otic epithelium and a subtle difference in otic *hmx3a* expression. **A**. Lateral view of otic vesicle and *dlx5a* expression in wild-type and *bmper* mutant embryos from 27 hpf to 72 hpf; anterior to the left. Black arrowheads indicate region where expression is absent in *bmper* mutants at 36 and 50 hpf. **B**. Analysis of *hmx3a* expression at 40 hpf and 50 hpf, lateral view of the otic vesicle, anterior to the left. Measurements were taken corresponding to the numbered white lines shown in the image of the wild-type ear at 40 hpf in B. 1. anterior gap; 2, anterior expression; 3, dorsal gap in expression; 4, posterior expression. The data are shown in the graphs below with the corresponding number. Two-way ANOVA with Šidák’s correction for multiple comparisons: \*\*\**p*=0.0003; ns, not significant. Levels of otic *hmx3a* expression appear unaffected, but the dorsal gap in expression increases in size. The region affected overlaps with *bmper* and *dlx5a* expression. Scale bars: 50 μm in A for all images.

Despite the obvious structural defect in the ear, additional markers that we tested were expressed normally, including those for the otic placode (*sox10, pax2a*), patterning at the early otic vesicle stage (*aldh1a3, hmx2a, otx1b*), sensory patch development (*pax2a*) and otic neurogenesis (*neurod*) (Supplementary Fig. S4). Of note, the normal expression domain of *otx1b* in ventral otic epithelium correlates with the normal formation of the ventral pillar and lateral (horizontal) semicircular canal duct (Fig. 1). Other markers for dorsal otic epithelium (*dlx3a, gbx2, dacha, lmx1ba, fn1a*) and the endolymphatic duct (*foxi1*) were expressed normally in *bmper* mutants (Supplementary Fig. S4). At 30 hpf, *bmper* is strongly expressed at the anterior and posterior poles of the otic vesicle; however, three of the genes that are co-expressed at this time, *fn1a*, *prdm1* and the proliferation marker *pcna*, all had unaltered patterns of expression in *bmper^sa108^* mutant embryos (Supplementary Fig. S4). Similarly, we analysed other markers of semicircular canal morphogenesis required for the formation of the epithelial projections that form pillars of tissue that will become the internal walls of the semicircular canal ducts (*adgrg6, vcana, ugdh, lmo4b, lmx1bb*). All these markers were expressed at normal levels in mutant embryos (Supplementary Fig. S4).

### Effect of the *bmper^sa108^* mutation on BMP signalling in the developing ear

To examine the effect of the *bmper* mutation on BMP signalling, we examined the expression of genes coding for BMP signalling pathway components in and around the developing ear. We compared expression of *bmp2b, bmp4, bmp6, bmp7b*, *smad6a, smad6b* and *chordin* in wild-type and *bmper* mutant otic vesicles at 30 hpf, a stage just prior to the first morphological changes seen in *bmper* mutant embryos, and at 48 hpf, when subtle changes in otic morphology become evident. However, we did not detect any significant differences in expression of BMP pathway genes in *bmper* mutant ears (Fig. 5A).

**Figure 5.**
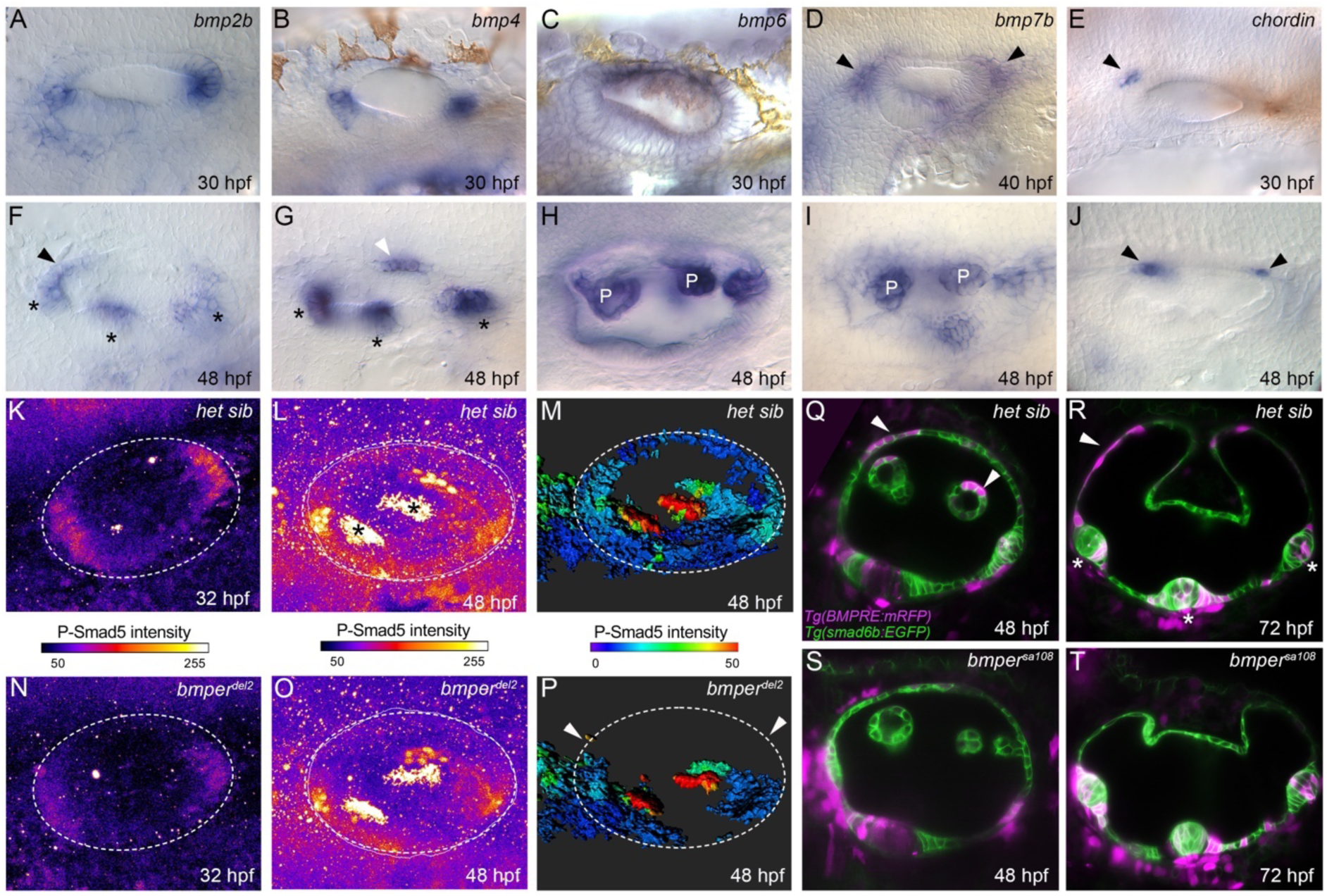
BMP pathway gene signalling is reduced in *bmper^sa108^* mutants. **A–J.** In situ hybridisations showing no change in the expression of BMP pathway genes in the otic vesicle at 30-40 hpf (A-E) and 48 hpf (F-J). Lateral views; anterior to the left. Expression is seen in the sensory cristae (*bmp2b, bmp4*, black asterisks, F,G); the lateral dorsal otic epithelium (*bmp2b*, black arrowhead, F); the ED (*bmp4*, white arrowhead, G);the non-sensory epithelial projections (*bmp6*, *bmp7b*, labelled P in H,I); and the periotic mesenchyme (*bmp7b*, *chordin*, black arrowheads, D,E,J). **K-P**. P-Smad5 levels (antibody stain), indicating BMP activity in the otic vesicle at 32 hpf (K,N) and 48 hpf (L,O) in wt (K,L) and MZ *bmper^del2^* mutants, lateral view, anterior left. A reduction of activity is seen in the MZ *bmper^del2^* otic vesicle at the anterior and posterior poles at 32 hpf. High levels of activity are seen in the sensory maculae at 48 hpf in heterozygous siblings and *bmper^del2^* mutants (black asterisks, L). M,P. Relative P-Smad5 levels at 48 hpf show a reduction of P-Smad5 staining in the dorsal otic epithelium (white arrowheads) and a small gap in the ventral otic vesicle (white asterisk). **Q–T.** Expression of the *Tg(BMPRE:mRFP)* reporter line in the *Tg(smad6b:EGFP)* background. mRFP fluorescence corresponding to BMP activity can be seen in phenotypically wild-type embryos in the dorsal otic vesicle at 48 hpf (Q) and 72 hpf (R) and also in the epithelial projections at 48 hpf (white arrowheads) Sensory cristae are shown with white asterisk (R). This expression in non-sensory epithelium is absent in *bmper^sa108^* mutant embryos (S,T). Expression in the cristae is unaffected in the mutant ear; expression in periotic mesenchyme appears up-regulated.

At 30 hpf, expression of *bmp2b* and *bmp4* is strong in the anterior posterior poles of the otic vesicle, overlapping with expression of *bmper*, whereas *bmp6* is only expressed dorsally. *bmp7b* is expressed in the periotic mesenchyme in two dorsal areas at the anterior and posterior of the vesicle adjacent to the region of *dlx5*a expression that is absent in *bmper* mutants (Fig. 5A). By 48hpf, *bmp2b* and *bmp4* expression expands dorsally; both still overlap with *bmper* but in different regions. Expression of *bmp4* overlaps with that of *gbx2* at the site of the developing ED, and *bmp2b* has weak expression in the anterior dorsal region, where *dlx5a* expression is lost. Both *bmp2b* and *dlx5a* are also strongly expressed in the cristae. *bmp6* and *bmp7b* become strongly expressed in the epithelial projections at 48 hpf. Two genes coding for negative regulators of BMP signalling are also expressed at these stages. We observed a transient focus of expression of *chordin* in the dorsal periotic mesenchyme, overlapping with the expression of *bmp7b* at 30 hpf. Subtle increases in the area of *bmp7b* expression domain in the periotic mesenchyme were noted in *bmper* mutants. The inhibitory SMAD gene *smad6b* is strongly expressed throughout the otic vesicle (Supplementary Fig. S2). However, we found no change in levels of GFP fluorescence in *bmper* mutant ears, or in the strength of the *bmper* otic phenotype, in the *Tg(smad6b:EGFP)* background.

To determine if BMP signalling activity was affected in the *bmper* mutant ear, we analysed levels of P-Smad5 in the otic vesicle at 32 hpf and 48 hpf using quantitative immunofluorescence. The strongest activity at 32 hpf was seen at the anterior and posterior poles of the otic vesicle, where *bmp2b* and *bmp4* are also expressed. In *bmper^del2^* mutant embryos, P-Smad5 levels were significantly decreased compared with heterozygous siblings. By 48 hpf, the strongest BMP activity was detected in the two sensory maculae, with low levels of P-Smad5 elsewhere in the otic epithelium. By comparing relative levels of P-Smad5, a reduction in BMP pathway activity was detected in non-sensory dorsal otic epithelium of *bmper^del2^* mutant embryos (Fig. 5K–P).

To confirm this result and measure BMP pathway activity in live embryos, we crossed *bmper^sa108+/-^* fish to the BMP reporter line *Tg(BMPRE:mRFP)* (Reichert et al., 2013). From 24 hpf, mRFP expression was predominantly in the anterior and posterior poles of the otic vesicle and also in the lateral region where the lateral projection will form. At 24 hpf, no difference was detected between wild-type and *bmper* mutant ears; however, the mRFP is not destabilised and may persist from earlier expression. A clear difference between wild-type siblings and *bmper* mutants was seen by 48 hpf: in wild-type embryos, mosaic expression of mRFP was observed in dorsal otic epithelium, which was absent in the *bmper* mutant ear. At 72 hpf, the semicircular canal duct truncations were already evident in the *bmper* mutant ear; mRFP expression was absent in canal wall epithelium in mutant embryos, but was strongly expressed in the cristae in both the wild-type and *bmper* mutant ear (Fig. 5Q–T). Taken together, the reduction in P-Smad5 staining and fluorescence levels of the BMP transgenic reporter suggest that Bmper is normally required to promote BMP signalling in in non-sensory otic epithelium, but does not appear to be required for BMP signalling in otic sensory tissue. We also noted a slight increase in mRFP fluorescence levels surrounding the otic vesicle at both stages in *bmper* mutants (Fig. 5Q–T). This might indicate that Bmper is required to repress BMP pathway activity in periotic mesenchyme.

### There are no substantial neural crest, cardiovascular or pronephric abnormalities in *bmper* mutant embryos

Over-expression studies in the chick and morpholino-based knockdown in the zebrafish have implicated Bmper in promoting development of the neural crest (Coles et al., 2004; Reichert et al., 2013). To test whether neural crest fates are affected in the *bmper^sa108^* mutant, we examined development of neural crest derivatives (craniofacial cartilages, pigment cells and Schwann cells). Although the cartilage stains were more variable in mutants, all elements of the craniofacial skeleton were present (Fig. 6). Pigmentation of larval and adult fish appeared largely normal; all three pigment cell types (melanocytes, xanthophores and iridophores) were present, and the stripe pattern was normal (Fig. 6). In addition, expression of the neural crest markers *foxd3* and *sox10* was present and apparently unaffected in *bmper^sa108^* mutant embryos, as was expression of *myelin basic protein* mRNA marking the presence of neural-crest-derived Schwann cells (Fig. 6).

**Figure 6.**
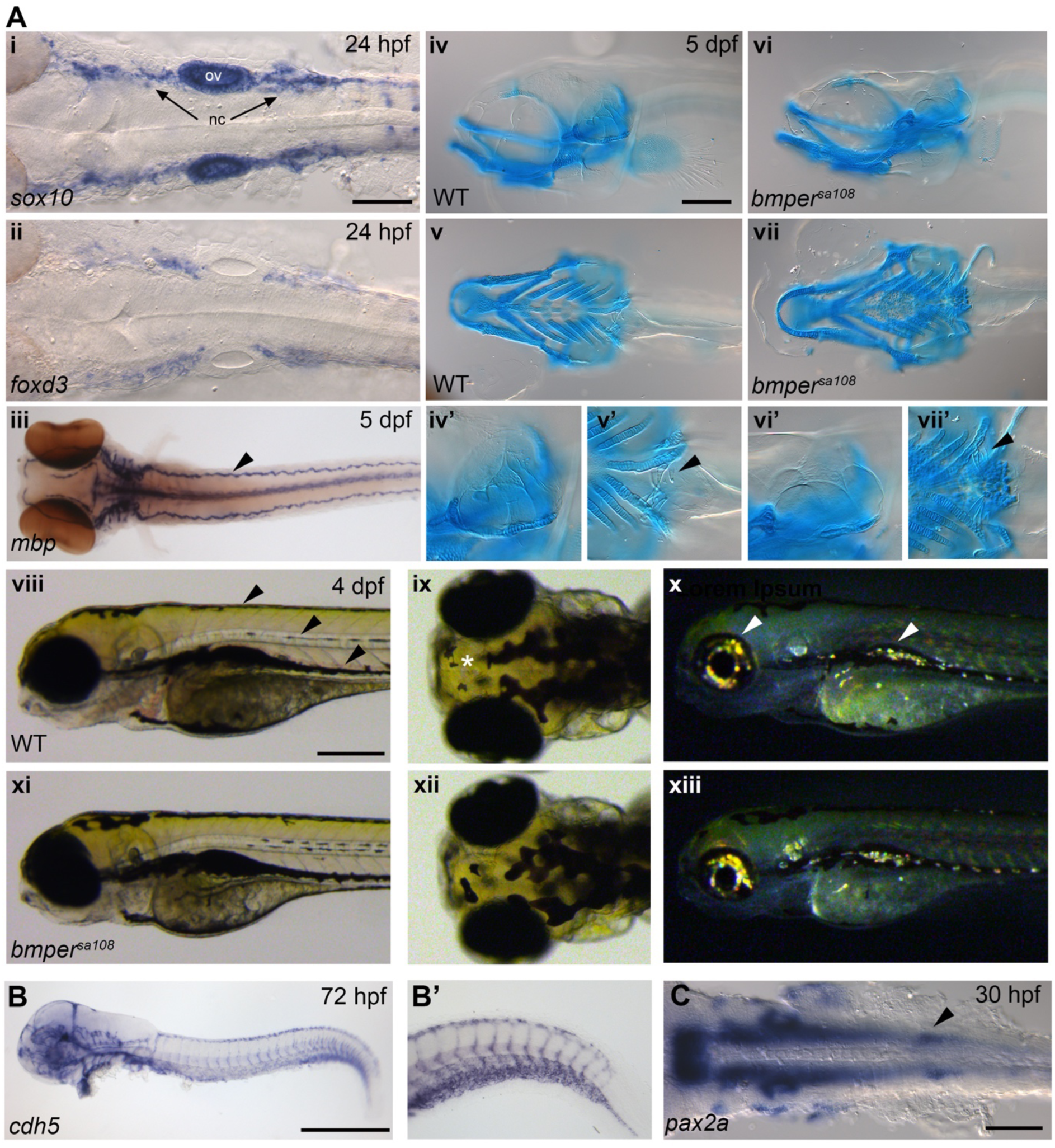
*bmper^sa108^* mutants do not display neural crest, angiogenesis or kidney development defects. **A. i-ii** Dorsal view of 24 hpf embryo showing expression of *sox10* (i) and *foxd3 (*ii). (iii) Dorsal view of 5 dpf larvae showing normal expression of *mbp*. The arrowhead marks expression along the posterior lateral line nerve. (iv-viii) Alcian blue staining of cartilage in wild-type (iv, v) and *bmper* mutant (vi, vii), lateral view (iv, vi) dorsal view (v, vii) iv’-vii’ detailed view of panels above showing the structure of the ear in wild-type and *bmper* mutants (black arrowhead)(iv’, vi’) and the presence of pharyngeal teeth in both wild-type and *bmper* mutant (v’, vii’)(arrowhead). (viii-xiii) pigmentation is normal in *bmper* mutants (xi-xiii) as in wild-type, including the normal patterning of three rows of melanocytes in the trunk (viii, arrowheads), xanthophores (yellow, white asterisk) (ix, xii); and iridophores (white arrowheads) (x, xiii). **B.** Lateral view of *ve-cadherin* expression at 72 hpf. Expression is normal in *bmper* mutants; see detail in the tail (B’). **C.** *pax2a* expression in the developing pronephros (black arrowhead) at 30 hpf is normal. Abbreviations ov, otic vesicle; nc, neural crest. Scale bars: 100 μm in Ai, for Aii, 200 μm in Aiv, for Av, vi, vii; 500 μm in Aviii, for Ax, xi, xiii, 500 μm in B; 200 μm in C.

Previous studies employing morpholino-mediated knockdown of *bmper* have reported defects in the tail vasculature, including defects in the intersegmental vessels (ISVs), dorsal longitudinal anastomotic vessel (DLAV) and caudal vein plexus (CVP) (Dyer et al., 2015; Heinke et al., 2013). To check whether there were any similar abnormalities in *bmper^sa108^* mutants, we examined the expression of *ve-cadherin* mRNA, a marker of vascular endothelium. We did not find similar defects or any consistent changes in vasculature patterning in developing *bmper* mutants (Fig. 5). Early patterning of the pronephros was assessed by *pax2a* expression, also unchanged in *bmper* mutants. Taken together, our analyses indicate that there are no substantial defects in the development of neural crest or cardiovascular derivatives in zygotic *bmper* embryos in the zebrafish, although we cannot rule out subtle defects in these or other tissues.

Previous studies have described moderate dorsalisation phenotypes following morpholino-mediated knockdown of *bmper* in the zebrafish, indicating an early requirement in promoting BMP signalling to establish ventral identity in the early embryo (Moser et al., 2007; Rentzsch et al., 2006). We have not found any evidence for this in the zygotic *bmper* mutants; embryonic axial patterning appeared normal in all embryos.

### Anatomy of the ear in adult *bmper^sa108^* mutants

The zebrafish zygotic *bmper* mutant alleles described here are adult viable and fertile. To examine the effect of the embryonic otic patterning defects on post-embryonic ear development we used lightsheet microscopy to analyse ear morphology at 21 dpf. At this stage in both wild-type and *bmper* mutant ears, the semicircular canal ducts have begun to elongate and the truncated canal ducts in the mutant have separated from the dorsal tissue.

In some specimens, a section of the canal duct remains attached to the crus commune, which is clearly visible. In the *bmper* mutant there can be a significant difference in the length of the anterior and posterior canals. The 90° angle between the anterior and posterior canals can be seen in the wild type and in *bmper* mutant ears. In all juvenile and adult ears that we examined, the lateral (horizontal) canal was present, complete, and relatively normal in morphology. All three ampullae, each containing a crista, were present.

To study the adult ear anatomy, we dissected ears from fixed adult homozygous *bmper^sa108^* mutant fish. Truncations of the anterior and posterior semicircular canal ducts persist into adulthood (Fig. 7). Some ears displayed severe truncations, where the canal duct was no more than a small bud attached to the ampulla (Fig. 7A-C), but in others the canal ducts extended some distance from the ampulla before ending blindly (Fig. 7E). The *Tg*(*smad6b:EGFP*) transgene is still expressed in adult ear tissue, enabling the cellular and structural detail of the adult canals to be imaged using this fluorescent marker. The truncated semicircular canal ducts have a normal diameter and curve in the correct orientation compared to the wild type.

**Figure 7.**
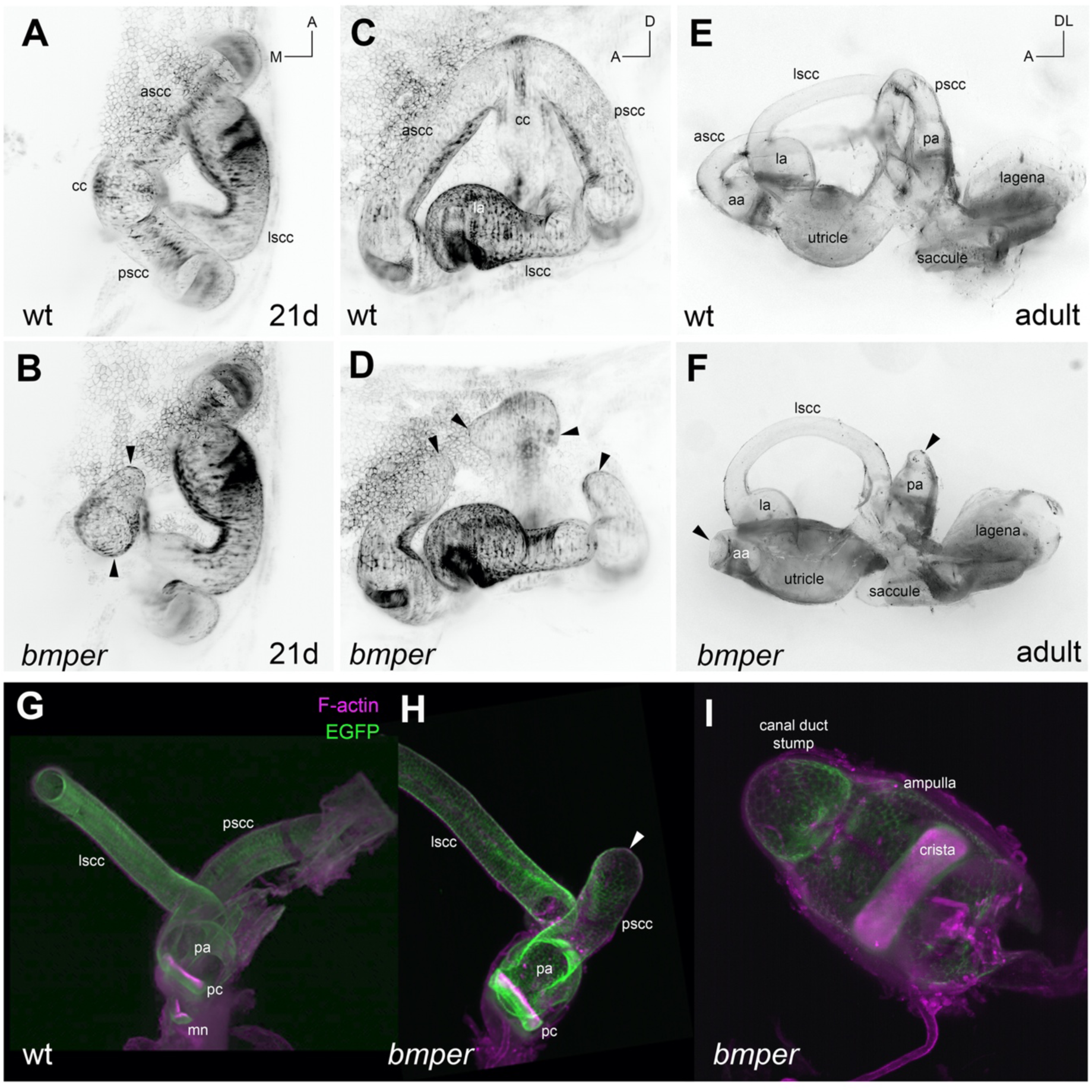
Truncations in the anterior and posterior semicircular canals persist into adulthood in *bmper* mutants. **A-D.** Day 21 wild-type (A, C) and *bmper* mutant (B, D) ears expressing the transgene *Tg(smad6b:EGFP*) and imaged using light-sheet microscopy. A,B Dorsal view, anterior to the top, medial to the left. C,D, lateral view, anterior left. **E,F.** Brightfield image of a dissected adult ear from wild-type (E) and *bmper* mutant (F), both dorsal view. Note truncated canals (black arrowhead). The lateral canal is present in *bmper* mutants (F). **G-I.** Fluorescent image of adult *Tg(smad6b:EGFP)* canals, counterstained with TRITC-phalloidin, dissected from wild-type and *bmper* mutant adults. H. Dissected canals from adult *bmper* mutant with a longer truncated posterior canal, compared with example in I (enlarged) where the canal terminated just above the ampulla. Arrowheads mark canal duct truncations. Abbreviations: A, anterior; D, dorsal; DL, dorsolateral; M, medial; cc, crus commune; aa, la, pa, anterior, lateral and posterior ampullae; pc, posterior crista; ascc, lscc, pscc, anterior, lateral and posterior semicircular canal ducts; mn, macula neglecta (not included in dissection in H).

## Discussion

### Zygotic Bmper function is required for correct morphogenesis of the anterior and posterior semicircular canal ducts in the developing ear

We have characterised two new mutant alleles of the BMP regulator gene *bmper*, both of which cause dorsal truncations of the anterior and posterior semicircular canals in the zebrafish inner ear. The nature and specificity of the ear phenotype were unexpected; to our knowledge, otic defects have not been described in murine *Bmper* mutants, and previous studies based on morpholino-mediated knockdown and mRNA mis-expression in the zebrafish have inferred roles for Bmper in embryonic axial patterning, placodal and neural crest specification and endothelial development, phenotypes that are not recapitulated in the zygotic mutants. However, the otic defects we describe in the zebrafish *bmper* mutants fit well with the known conserved role of BMP signalling in the inner ear.

BMP signalling is known to play an important role in semicircular canal formation in mice, chick and zebrafish (reviewed in (Alsina and Whitfield, 2017)). Our previous work demonstrated a requirement for Bmp2b function in the development of all three semicircular canal ducts in the zebrafish ear (Hammond et al., 2009), and this requirement is conserved in mice (Hwang et al., 2019). The *bmper* mutant phenotype is more specific and less severe than that in *bmp2b* mutants, only affecting the anterior and posterior ducts, and resulting in dorsal truncations of these structures in the adult ear. In amniote ears, the anterior and posterior canal ducts, together with the crus commune, develop from a dorsolateral pouch, whereas the horizontal (lateral) canal develops from a lateral pouch (reviewed in (Alsina and Whitfield, 2017)). Thinning of the dorsolateral pouch in the chick is dependent on BMP signalling, and correlates with pSMAD activity (Ohta et al., 2010). Pouches are not so evident in the developing zebrafish ear, but the *bmper* mutant phenotype demonstrates that the anterior and posterior canals share genetic requirements that are separate from those for the lateral (horizontal) canal, as in amniotes. We therefore propose that a region of the zebrafish otic vesicle exists that is the equivalent of the dorsolateral canal pouch in amniote ears, and is dependent on the promotion of BMP signalling by Bmper for its normal morphogenesis.

The lateral (horizontal) canal duct develops relatively normally and is complete in the adult *bmper* mutant ear. This reflects the normal expression of *otx1b* at otic vesicle stages in *bmper* mutants, and normal appearance of the ventral pillar. Otx1 function is known to be required for formation of the lateral canal in both zebrafish and mammals (reviewed in (Alsina and Whitfield, 2017)). The endolymphatic duct, which is a dorsal otic structure, also appears to be unaffected in the *bmper* mutant ear. Interestingly, endolymphatic duct outgrowth was also unaffected by overexpression of the BMP inhibitor gene Noggin in the chick (Ohta et al., 2010). Taken together, our results suggest that patterning of the otic dorsoventral axis is unaffected in *bmper* mutants, but reveal a specific requirement for Bmper in the morphogenetic events that convert dorsolateral otic epithelium into the anterior and posterior canal ducts.

It is interesting that all three ampullae and cristae apparently form normally in the *bmper* mutant ear. Expression of BMP genes in cristae is highly conserved across vertebrates (Hemmati-Brivanlou and Thomsen, 1995; Morsli et al., 1998; Mowbray et al., 2001; Wu and Oh, 1996), and crista development is dependent on Bmp4 function in the mouse (Chang et al., 2008). Moreover, mis-expression of inhibitors of BMP signalling (Smad6, Noggin) resulted in the down-regulation of expression of crista-associated genes in the chick (Chang et al., 2008). However, we found that ampullae and cristae formed in the absence of Bmp2b function in the zebrafish (Hammond et al., 2009), as in the *bmper* mutants. The *bmper* gene has an expression domain that overlaps with that of *bmp2b* in the zebrafish ear (Mowbray et al., 2001). These data may indicate that Bmper acts to modify Bmp2b-dependent signalling in the ear. Additionally, other BMPs known to interact with Bmper are co-expressed with *bmper* in different domains of the ear and may contribute to the defect in canal formation. These domains include the area of expression of *bmp4* and *bmp2b* at the anterior and posterior poles of the otic vesicle at 30 hpf, the expression of *bmp6* and *bmp7b* in the epithelial projections, and the expression of *bmp7b* and *chordin* in periotic mesenchyme adjacent to the site where the canal truncations form.

### Transcriptional targets of *bmper* expression in the zebrafish ear

Our results identify *dlx5a* as a likely transcriptional target of canonical BMP/SMAD signalling in the zebrafish ear, potentiated by Bmper. The *dlx5a* gene is expressed in a similar pattern to both *bmp2b* and *bmper* in the dorsal otic vesicle in zebrafish at the 20-25 somite stage (Solomon et al., 2003; Thisse and Thisse, 2004), and *dlx5a* expression is lost in dorsal otic epithelium (excepting the endolymphatic duct) in the *bmper^sa108^* mutant. A reciprocal relationship is found in the mouse, where several lines of evidence identify *Bmper* as a likely direct transcriptional target of Dlx5 in the otocyst (Sajan et al., 2011); murine *Bmper* expression is also dependent on Dlx5/6 in the branchial arches (Jeong et al., 2008).

*Dlx* genes have a known function in semicircular canal development in the mouse: canal ducts fail to form correctly in the murine *Dlx5* mutant ear (Acampora et al., 1999; Merlo et al., 2002), while in the double *Dlx5/6* mouse mutant ear, there is a loss of all dorsal derivatives, including the semicircular canals, correlating with a loss of otic *Gbx2*, *Bmp4* and *Otx1* expression at early stages (Robledo and Lufkin, 2006). It is likely that these changes are mediated, at least in part, through regulation of BMP signalling, as manipulation of BMP signalling results in similar phenotypes. Conditional disruption of Bmp4 in the mouse ear results in down-regulation of *Dlx5* expression, although *Dlx5* expression remains in the ED (Chang et al., 2008), similar to the pattern we see in zebrafish *bmper* mutants. Dorsomedial Dlx5 and Gbx2 expression is also responsive to Wnt signalling (Lin et al., 2005; Merlo et al., 2002).

Non-canonical BMP/PKA signalling also regulates the expression of *Hmx3* in the chick ear (Ohta et al., 2016). Our data revealed only subtle changes in the spatial extent of otic *hmx3a* expression in the *bmper* mutant ear, and *hmx3a* expression persisted in the dorsolateral regions where *dlx5a* expression is lost in *bmper* mutants. Thus, *hmx3a* does not appear to be a major transcriptional target of *bmper*-mediated BMP signalling in the zebrafish ear. Otic expression of *hmx3a* is instead known to be regulated by Fgf and Hedgehog signalling in the zebrafish, and is required for normal otic anterior patterning (Hartwell et al., 2019).

### Structure and function of the zebrafish Bmper protein

Bmper, together with its *Drosophila* orthologue Crossveinless-2 (Cv-2), is a known regulator of BMP signalling with both pro-BMP and anti-BMP activity in different contexts (Conley et al., 2000; Rentzsch et al., 2006; Serpe et al., 2008; Zhang et al., 2010). In the fly, secreted Cv-2 protein interacts with heparan sulphate proteoglycans on the cell surface, where it fine-tunes BMP signalling in a concentration-dependent manner (O’Connor et al., 2006; Serpe et al., 2008). In the fish, morpholino-mediated knockdown of *bmper* has been reported to cause moderate embryo dorsalisation, indicating that the protein normally functions to promote BMP activity during patterning of the embryonic dorsoventral axis (Moser et al., 2007; Rentzsch et al., 2006; Zhang et al., 2010). However, injection of full-length *bmper* mRNA gave rise to a full spectrum of dorsalised and ventralised phenotypes, suggesting both pro- and anti-BMP activity (Rentzsch et al., 2006; Zhang et al., 2010). The proposed explanation for this apparently contradictory finding is that cleavage of Bmper converts it from an anti- to a pro-BMP factor: injection of mRNA coding for an uncleavable protein has strong dorsalising activity, while that coding for the cleaved N-terminal fragment alone (amino acids 1-355) has strong ventralising activity (Rentzsch et al., 2006). Zhang and colleagues report that the anti-BMP function is effected by binding to BMP, while the pro-BMP function is mediated through binding to Chordin (Zhang et al., 2010). The different cleaved versions are also proposed to have similar affinities to BMPs but different affinities to the extracellular matrix which is thought to interact with Bmper through the C terminus (Rentzsch et al., 2006). The data we have presented here for the zebrafish *bmper* mutant phenotype, including the comparison to known BMP loss-of-function otic phenotypes and the reduction in P-Smad5 and BRE activity in otic tissue, all indicate a pro-BMP function for Bmper in the zebrafish ear.

The Bmper protein has structural similarity with the BMP inhibitor protein Chordin (Rentzsch et al., 2006), and some conservation with the larger glycoproteins alpha-Tectorin and Otogelin, which have roles in otolith tethering in the developing zebrafish ear (Stooke-Vaughan et al., 2015). However, otolith formation and tethering appear normal in the *bmper* mutant ear (this work), and semicircular canals form normally in *tecta* and *otog* mutants (Stooke-Vaughan et al., 2015), so the functions of these structurally related glycoproteins do not appear to overlap in the zebrafish ear.

It is likely that the discrepancies between the mutant phenotype described here and the much more severe effects on embryonic patterning due to morpholino knockdown (Moser et al., 2007; Rentzsch et al., 2006; Zhang et al., 2010) are partly due to genetic compensation via transcriptional adaptation (El-Brolosy et al., 2019; Sztal and Stainier, 2020). However, it is also possible that the *bmper^sa108^* allele is hypomorphic, and that the mutant protein, if produced, retains some function. The STOP codon is downstream of the cleavage site after amino acid 353, and so the mutant protein might be expected to undergo cleavage and generate an N-terminal fragment with pro-BMP (ventralising) function. The ear phenotype, however, is indistinguishable between the *bmper^sa108^* and *bmper^del2^* alleles, and thus both may represent complete loss of function for otic morphogenesis. Bmper has been shown to bind Bmp2/7 through vWFC1 (Zhang et al., 2007), which remains intact in the predicted mutant proteins, and so it is possible that some anti-BMP function is retained in the mutants, and is perhaps active in other tissues.

### Zebrafish Bmper does not appear to function in establishment of the preplacodal region, otic placode or neural crest

Competence to form the pre-placodal region (PPR) that gives rise to the otic placode relies on active BMP signalling at late blastula stages, followed by a specification step involving attenuation of BMP signalling in the late gastrula (Bhat et al., 2013) (and references within). Previous studies using morpholino-based knockdown have suggested that Bmper acts as a BMP antagonist in generation of the PPR. Kwon and Riley proposed that *bmper* is one of a number of BMP antagonist genes expressed in cephalic paraxial mesendoderm that together contribute positively to otic induction from the PPR, although they found ‘little or no effect’ on development of the ear following morpholino-mediated knockdown of *bmper* alone (Kwon and Riley, 2009). On the other hand, Esterberg and Fritz reported expression of *bmpe*r in the PPR (ectoderm) itself, and that morpholino-mediated knockdown of *bmper* resulted in a circular otic vesicle with no otoliths, similar to that of *dlx3b/4b* morpholino knockdown (Esterberg and Fritz, 2009). Using morpholino-based knockdown and mRNA mis-expression experiments, these authors also suggested a role for Bmper in attenuating BMP activity to establish the PPR and to confer competence to respond to FGF signalling (Esterberg and Fritz, 2009). However, a subsequent study has challenged this interpretation; Reichert and colleagues have shown that *bmper* is expressed in the zones of higher BMP activity either side of the PPR, and propose that it acts locally to promote BMP activity and expression of neural crest markers. Their study identifies the BMP antagonist *bambib* as responsible for attenuation of BMP signalling in the PPR (Reichert et al., 2013).

In zygotic homozygous *bmper* mutants, we see very little effect on PPR specification, otic placode induction, early otic vesicle shape, otolith formation or neural crest specification. Notably, we do not find any reduction in expression of the otic placode marker *pax2a*, or of the neural crest markers *foxd3* and *sox10*, contrary to previous reports of down-regulation of these genes following morpholino-based knockdown of *bmper* (Esterberg and Fritz, 2009; Reichert et al., 2013). If *bmper* conferred competence to respond to FGF signalling during otic induction and patterning, we might expect the mutant otic phenotype to resemble that of embryos with compromised FGF signalling (Hammond and Whitfield, 2011; Hartwell et al., 2019; Léger and Brand, 2002; Maier and Whitfield, 2014)—but this is not the case. Our results, therefore, do not support a role for Bmper in PPR specification, otic placode induction or neural crest specification.

In the mouse, a differential RNAseq study of dissected otocysts identified *Bmper* as a significantly upregulated transcript following conditional ablation of *FGFR2b* over different time windows between E9 and E12; ventral expansion of the *Bmper* otic expression domain was validated by in situ hybridisation (Urness et al., 2018). These results implicate signalling via the FGFR2b ligands FGF3 and FGF10 in limiting *Bmper* expression to dorsal otic epithelium in the mouse.

### Expression and roles in other tissues, and comparison with mammalian *Bmper* mutant phenotypes

It is puzzling that the phenotype in the zebrafish *bmper* mutants appears to be so restricted to the ear. The zebrafish *bmper* gene itself is expressed in many other tissues, including paraxial mesendoderm, neural plate border, epiphysis, lens, cloaca, and elements of the developing craniofacial skeleton and vasculature (Clément et al., 2008; Esser et al., 2018; Esterberg and Fritz, 2009; Kessels et al., 2014; Kudoh et al., 2001; Kwon and Riley, 2009; Li et al., 2009; Lockhart-Cairns et al., 2019; Moser et al., 2007; Reichert et al., 2013; Rentzsch et al., 2006). It was not our intention here to provide a comprehensive analysis of tissues other than the ear, and it is quite possible that additional defects are present. However, the overall phenotype is clearly less severe in fish than in mammals (see below). This could reflect sub-functionalisation with a paralogous gene in the fish, but the *bmper* gene appears to be unique in the zebrafish genome; there is no predicted gene duplicate. Another possible interpretation is that the C-terminal truncations predicted for the *bmper^sa108^* and *bmper^del2^* alleles remove a portion of the protein that is critical for function in the ear, but not in other organ systems. It seems unlikely, however, that the severe truncation in the *bmper^del2^* allele is able to retain significant protein function.

Mutations in the murine *Bmper* gene cause cardiovascular, skeletal and renal phenotypes, all indicative of a pro-BMP function for the wild-type protein (Dyer et al., 2014; Ikeya et al., 2006; Ikeya et al., 2010). To our knowledge, the inner ear phenotype has not been characterised in the mouse, although *Bmper* is expressed in the murine otic vesicle (Coffinier et al., 2002; Ikeya et al., 2006; Urness et al., 2018). In humans, recessive mutations in *BMPER* are causative for a family of rare skeletal dysplasias including diaphanospondylodysostosis (DSD), symptoms of which include delayed or absent vertebral ossification, missing ribs and craniofacial and renal anomalies (Ben-Neriah et al., 2011; Funari et al., 2010; Gonzales et al., 2005; Greenbaum et al., 2019; Hofstaetter et al., 2018; Kuchinskaya et al., 2016; Legare et al., 2017; Zong et al., 2015) (OMIM 608022). The most severe cases are perinatal lethal due to respiratory insufficiency; hearing loss was reported in two patients surviving beyond infancy (Kuchinskaya et al., 2016; Scottoline et al., 2012). Four siblings presenting with severe congenital scoliosis, but a lack of renal anomalies, were shown to be transheterozygous for mutations in *BMPER* (Zong et al., 2015). Notably, two of the human mutations described predict a similar protein truncation to *bmper^sa108^*, with a loss of the C-terminal cysteine-rich and trypsin inhibitor domains (Kuchinskaya et al., 2016; Zong et al., 2015) (Fig. 3C). More recent studies have identified a role for BMPER in insulin resistance and obesity (Mao et al., 2021; Pérez et al., 2021), which could also be tested for in the adult viable zebrafish mutants. No study has been made of any vestibular anomalies in the human DSD patients, to our knowledge. The zebrafish mutants will be useful in prompting such studies, and may help to improve our understanding of the pathology of these rare disorders.

## Acknowledgements

We thank Caroline Hill for providing the *Tg(BMPRE:mRFP)* line, M. Angela Nieto for the *Tg(Xla.Eef1a1:h2b-mRFP1)* line and Henry Roehl for providing the *bmper* cDNA. We thank several members of the scientific community for helpful discussion, especially Patrick Blader, Caroline Hill and Nick Monk. We are grateful to the Sheffield aquarium staff for expert fish husbandry, and thank the Sanger Institute Zebrafish Mutation Resource for providing the *otx1b^sa96^* mutant allele, which led to the discovery of *bmper^sa108^* in the background in the Whitfield lab. The Mullins lab obtained the *bmper^sa108^* line directly from the Sanger collection.

## Funding

This work was funded by the BBSRC (BB/J003050 to TTW, BB/M01021X/1 and BB/S007008/1 to TTW and SB, and BB/D020433/1 to RDK). EM was supported by an EU Marie Skłodowska-Curie Intra European Fellowship (275978). Work in the Mullins lab was funded by NIH grants R35-GM131908 (MCM), T32-GM007229 and F31-GM113362 (FBT), T32-HD08318 and an NSF Graduate Fellowship (JZ). Imaging in Sheffield was carried out in the Wolfson Light Microscopy Facility, supported by a BBSRC ALERT14 award to TTW and SB for light-sheet microscopy (BB/M012522/1). The Sheffield zebrafish aquarium facilities were supported by the MRC (G0700091).

---

**Supplementary Figure S1.**
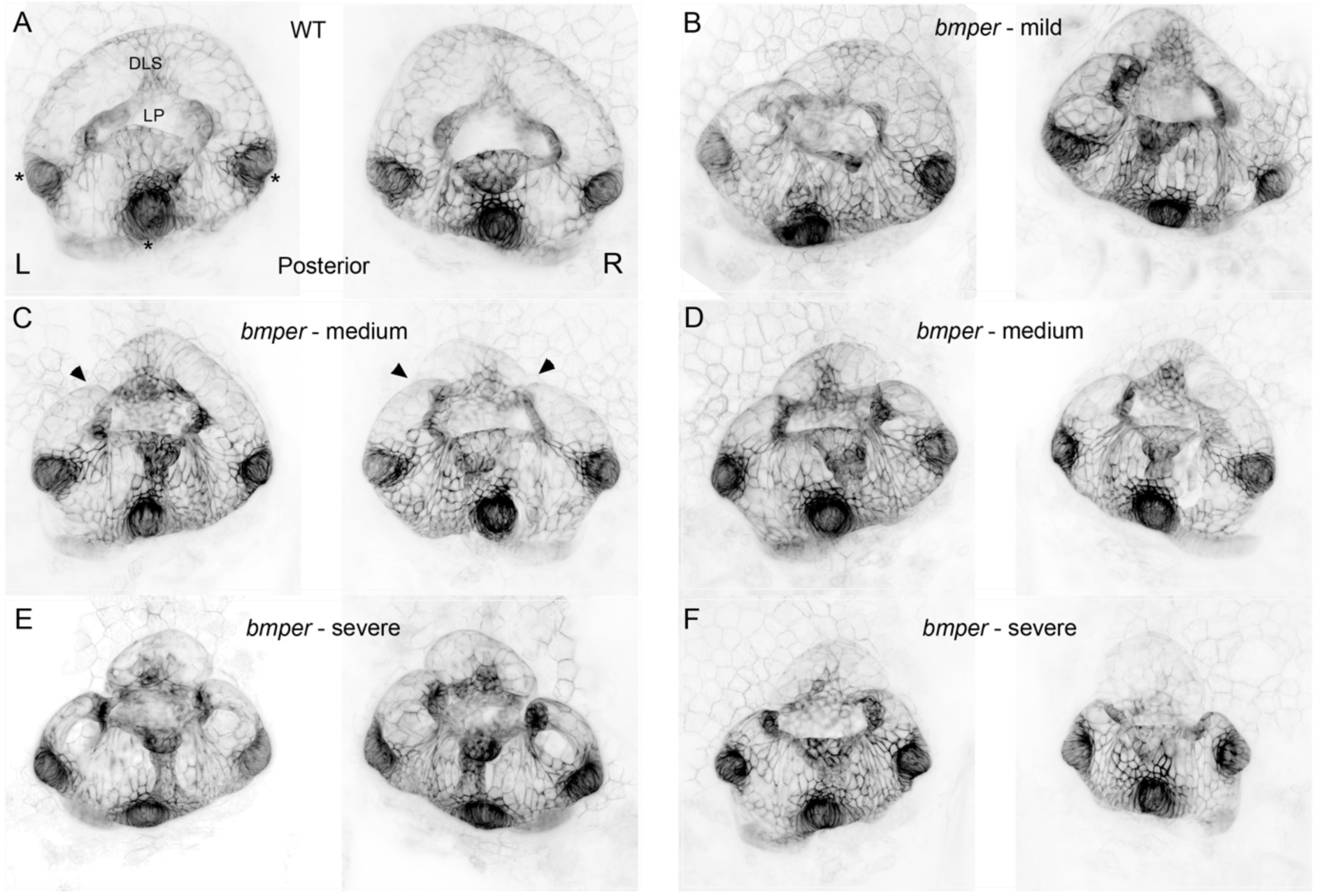
The *bmper* mutant phenotype results in variable levels of truncation in the anterior and posterior semicircular canal ducts. **A–F**. Light-sheet microscope inverted MIP images of the *Tg(smad6b:EGFP)* line, showing otic vesicles at 5 dpf. Each pair of panels show the left (L) and right (R) ear from the same embryo. The posterior canal is to the centre of each pair of panels. **A**. Wild-type ear showing the position of the DLS, lateral projection (LP) and the strong GFP expression in the cristae (black asterisks). **B–F.** Examples of ears from *bmper^sa108^* mutants. Variability can be seen between the different ears from the same embryo and also between different embryos. The embryo in B has a milder phenotype with the posterior canal intact on the left ear, compared with the severe phenotypes in D–F, with complete truncations of both semicircular canals in both ears. Truncations are highlighted in panel C (black arrowheads).

**Supplementary Figure S2.**
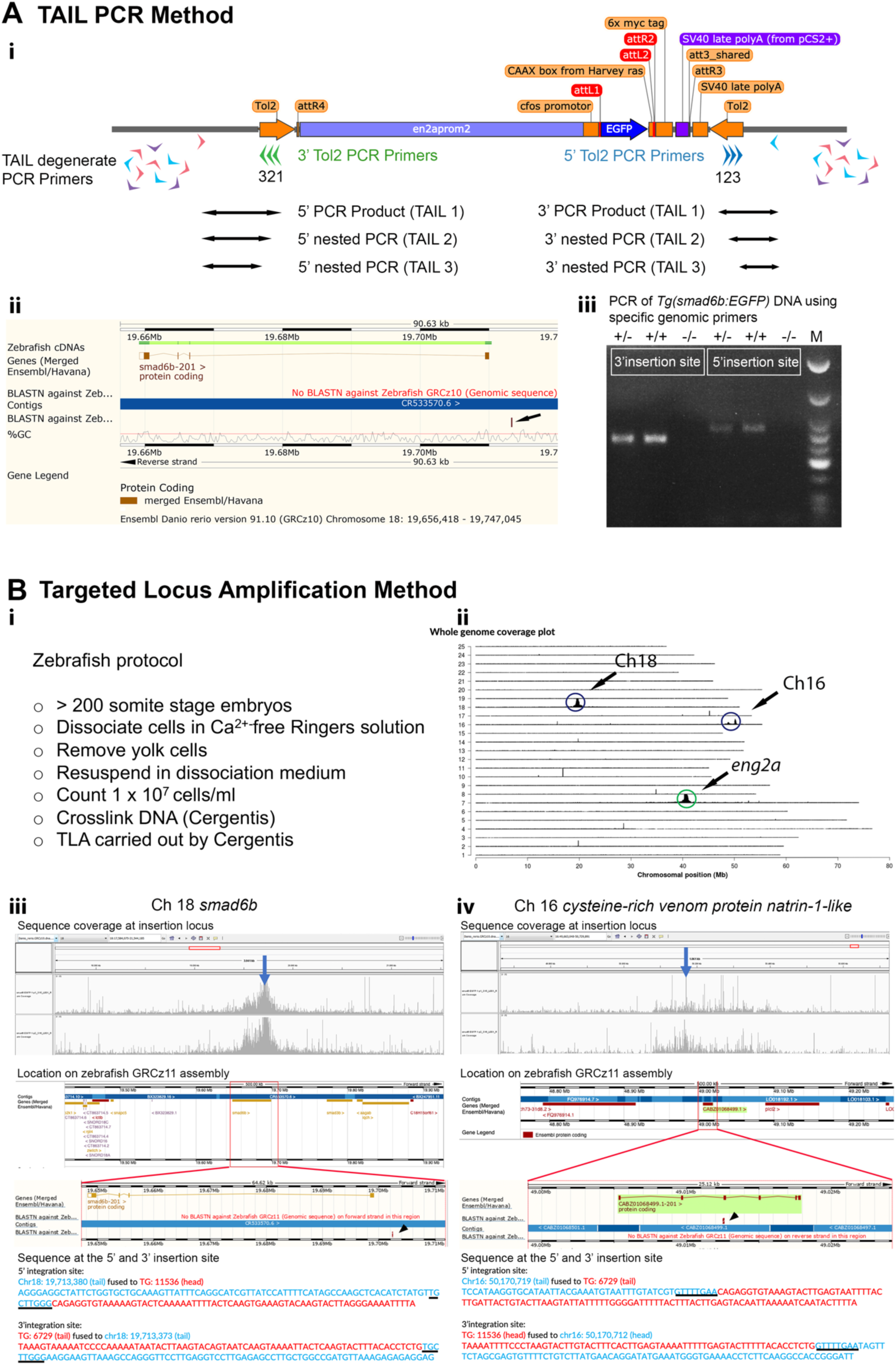
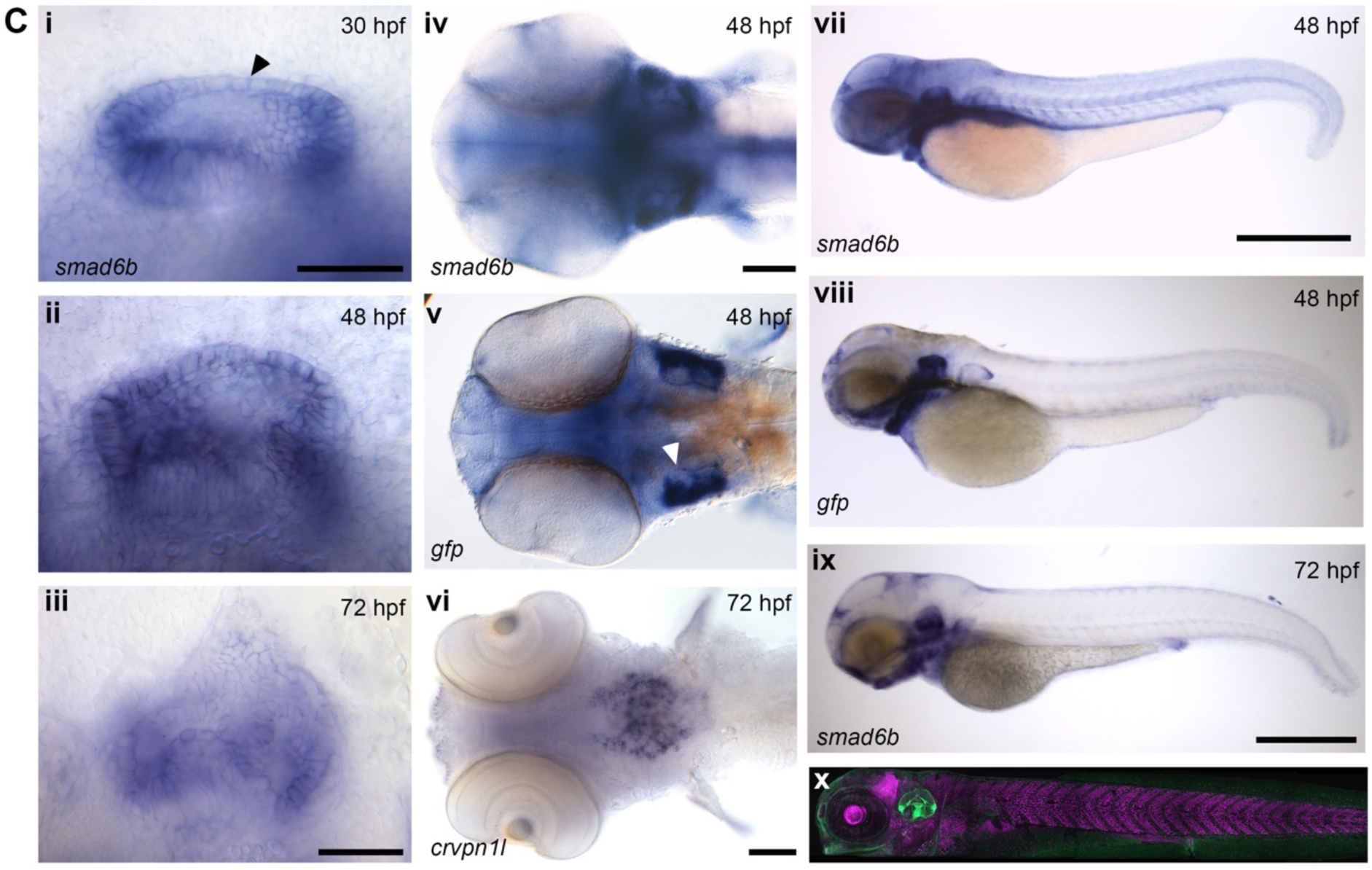
Identification of the genomic insertion site of a transgenic line expressing GFP in the inner ear. **A.** (i) Overview of the thermal asymmetric interlaced (TAIL-PCR) strategy. Schematic structure of the transgene, containing *engrailed2a* promoter sequences, a *cfos* minimal promoter and EGFP-CAAX coding sequence, flanked by Tol2 sequences. Tol2-specific primers are shown for the 5’ (green) and 3’ (blue) Tol2 sites. Degenerate primers are shown in red, blue and purple. PCR products from reactions using specific Tol2 primers and degenerate primers are shown. The products from nested PCR reactions (TAIL2 and TAIL3) were used to find flanking genomic sequences. ii PCR products from 5’ and 3’ reactions that map to the same position in the genome were found to match sequences at the 3’ end of the *smad6b* locus. iii The *smad6b* insertion site was confirmed using specific genomic primers and the Tol2-specific primers. **B.** (i) Protocol for preparation of zebrafish cell suspension for the Targeted Locus Amplification TLA method. (ii) Result of TLA sequence trace mapping to the zebrafish genome. Peaks can be seen at the same Ch18 locus close to *smad6b* found by TAIL PCR. In addition, another locus on Ch16 was found. The peak on Ch7 in the green circle corresponds to the endogenous *eng2a* locus. (iii) Detailed view of the sequences traces around the insertion site above an overview of the genomic location. Note *smad6b* is also close to *smad3b*. The insertion sequences map to the same 3’ region of *smad6b* in the zebrafish GRCz11 assembly. Below is the insertion sequences at the 3’ and 5’ insertion site. Note the target site duplication (TSD) 8 bp tandem repeated sequence, underlined in black. iv As in iii for the Ch16 insertion site. The insertion is in the intronic sequence of the *cysteine-rich venom protein natrin-1-like* gene. **C.** (i-iii) In situ hybridisation of *smad6b* expression throughout the otic vesicle at i, 30 hpf, ii, 48 hpf, iii, 72 hpf; lateral views, anterior to the left. (iv, v) Dorsal view at 48 hpf of *smad6b* expression (iv) and *gfp* expression (v), in *Tg(smad6b:EGFP)* embryos, anterior to the left. *gfp* and *smad6b* mRNA have closely matching spatial expression domains. Note the expression of *smad6* and *gfp* is weaker more medially and also more dorsally at 30 hpf (i). vi Ventral view of specific expression of *crvpn1l* at 72 hpf. vii, viii Whole mount in situ staining of 48 hpf embryos with *smad6b* (vii) and *gfp* (viii) *Tg*(*smad6b:EGFP*) embryos showing the overlap in expression. (ix), (x), 72 hpf *Tg*(*smad6b:gfp*) embryos showing strong expression of *smad6b* in the ear in (ix) and strong otic expression of GFP (green) in a confocal image (x). Nuclei are labelled with *Tg(Xla.Eef1a1.mRFP1)* (magenta). Scale bars: 50 μm in Ci, for Cii, Ciii; 100 μm in Civ, for Cv, Cvi; 500 μm in Cvii, for Cviii, Cix.

**Supplementary Figure S3.**
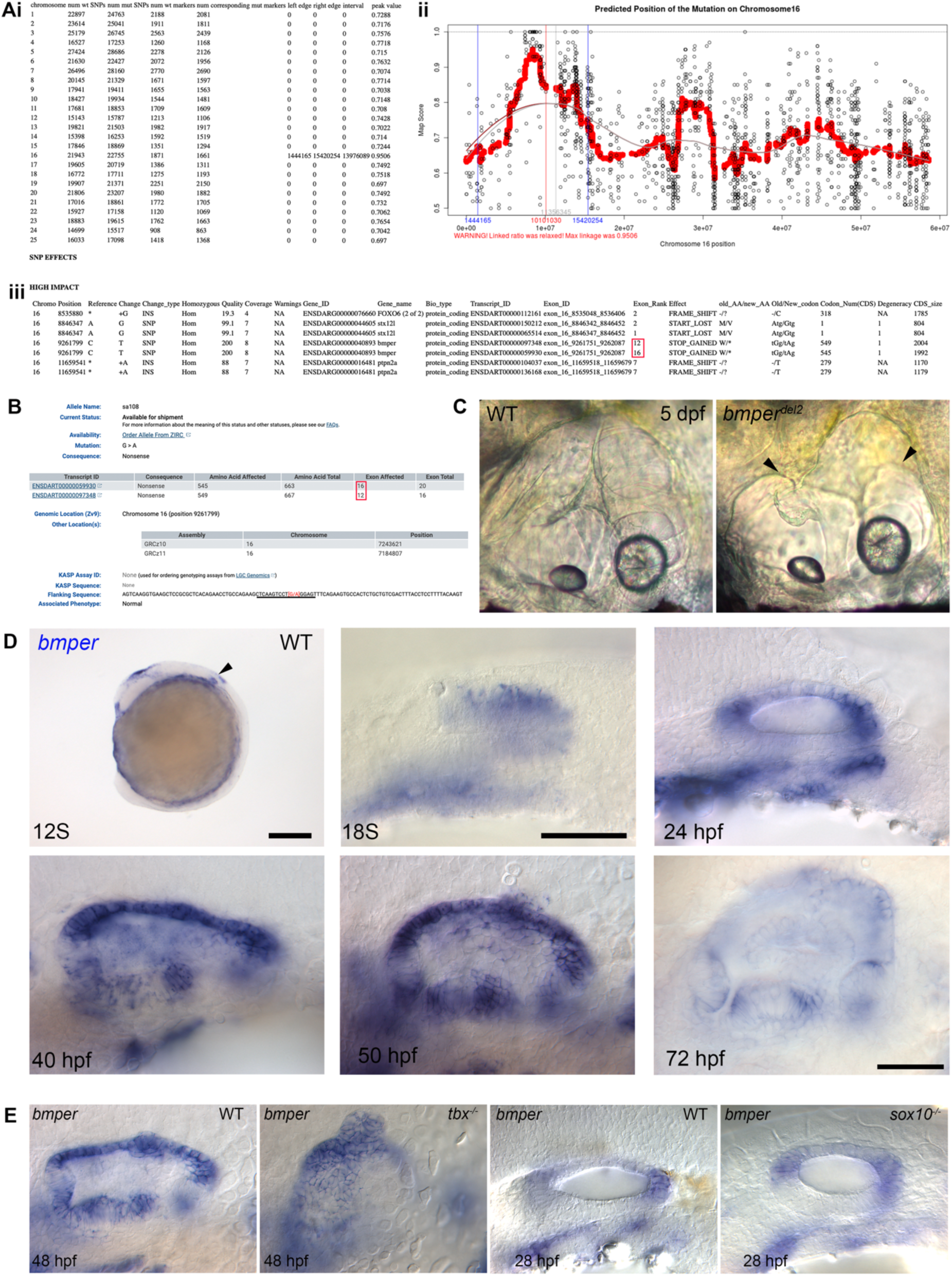
Independent confirmation of mutations in *bmper*. **A.** (i) RNAseq genome wide-data listing the numbers of wild-type and mutant markers per chromosome. A strong peak value (0.9506) representing a high mutant marker frequency is detected on Chromosome 16 at the interval 1444165-15420254. (ii) Graphical view of the mutant marker frequency at the candidate interval on Ch16. (iii) List of high impact SNPs and insertions occurring in genes within the candidate interval. A STOP is gained in the *bmper* gene, making this a strong candidate. **B.** Screenshot from the Sanger Institute Zebrafish Mutation Project website (sanger.ac.uk/resources/zebrafish/zmp) showing the *sa108* mutation in the *bmper* gene at the same amino acid residue as the mutation characterised in Fig. 3. **C.** DIC image of wild-type ear at 5 dpf and the ear in a *bmper^del2^* mutant, showing truncated anterior and posterior canal ducts (arrowheads). Compare to the *bmper^sa108^* mutant ear, Fig. 1A(vi). Lateral views, anterior to left. **D.** Expression of the *bmper* gene in the otic vesicle from 12 somites (12S) to 72 hpf. The strongest dorsal *bmper* expression is present just prior to the onset of epithelial projection outgrowth (42 hpf onwards). **E.** Expression of *bmper* in *tbx^-/-^* and *sox10^-/-^* mutant embryos at 48 hpf and 28 hpf, respectively. No difference in the level of expression was detected. Scale bars: 50 μm in D detailed view of ear; 200 μm in D, 12S whole mount.

**Supplementary Figure S4.**
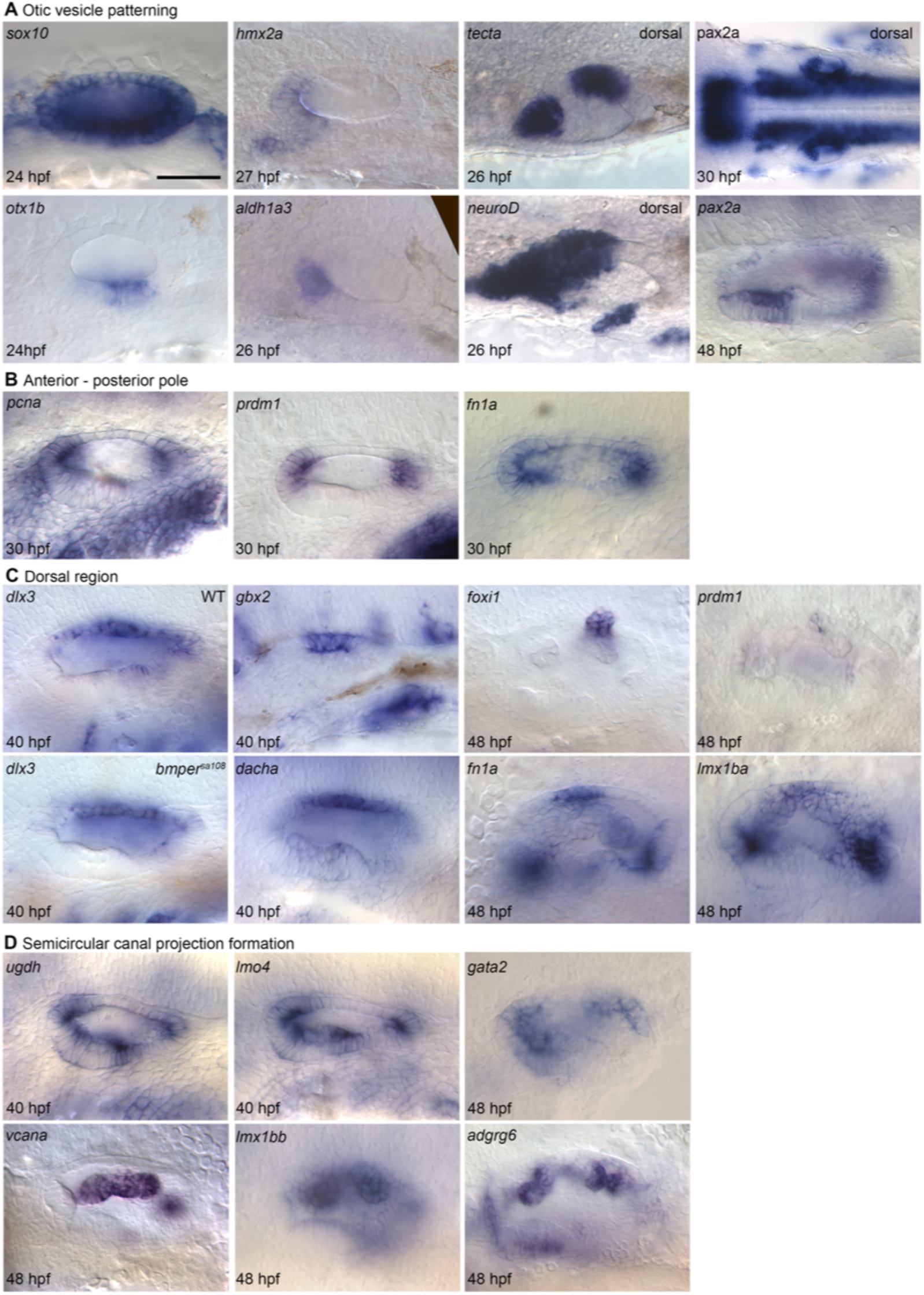
Otic patterning and dorsal gene expression that is unaffected in *bmper^sa108^* mutants. **A.** Otic vesicle patterning genes. **B** Genes strongly expressed in the anterior and posterior poles of the otic vesicle are unaffected in *bmper* mutants. **C.** Genes expressed in or around the endolymphatic duct. Note *dlx3* is unchanged in *bmper* mutants, unlike the premature down-regulation of the related gene *dlx5a* (Fig. 4A). **D.** Genes previously shown to be required for semicircular canal morphogenesis are also unaffected in *bmper* mutants. Scale bars: 50 μm in A for all images (except *pax2a* dorsal).

**Supplementary movie 1 – 3D rendered day 7 wild-type sibling ear**

3D rendered image of wt 7 dpf otic vesicle using light-sheet images of the *Tg*(*smad6b:EGFP*), line which marks cell membranes of the otic epithelium. The movie begins with a lateral view of the ear with posterior to the left, and as the ear rotates the medial view is seen. The external surface shows the three semicircular canals and the internal view shows the pillars that form the hubs for each canal, the three cristae and the septa that divide the canals.

**Supplementary movie 2 – 3D rendered day 7 *bmper* mutant ear**

3D rendered image of a *bmper* mutant 7 dpf otic vesicle using light-sheet images of the *Tg(smad6b:EGFP*) line. The movie begins with a lateral view of the ear with posterior to the left, and as the ear rotates the medial view is seen. The external surface shows the three semicircular canals, with a clear truncation in the anterior canal; the posterior canal appears normal. The internal view shows the pillars that form the hubs for each canal, the three cristae and the septa that divide the canals.

**Supplementary movie 3 – composite *z* stack movie of wild-type and *bmper* mutant otic vesicles from** **Figure 1**

Movie through the dorsal view of a wild type ear, mild *bmper* and stronger *bmper* phenotype. *z* slices from light-sheet microscopy of the *Tg(smad6b:EGFP)* line. The three embryos from Figure 1D are aligned so that the movie begins at an equivalent depth in each embryo and the semicircular canal truncations in the *bmper* mutants can be seen.

## Notes

### Competing Interest Statement

The authors have declared no competing interest.

