## Supplementary Table S1 for "Bmper is required for morphogenesis of the anterior and posterior semicircular canal ducts in the developing zebrafish inner ear"

**Supplementary Table S1. List of RNA in situ probes.**

| **Probe name** | **ZFIN ID** (zfin.org) | **Reference** |
|---|---|---|
| *adgrg6* | ZDB-GENE-041014-357 | (Geng et al., 2013) |
| *aldh1a3* | ZDB-GENE-061128-2 | (Pittlik et al., 2008) |
| *bmp2b* | ZDB-GENE-980526-474 | (Hammerschmidt et al., 1996) |
| *bmp4* | ZDB-GENE-980528-2059 | (Hammerschmidt et al., 1996) |
| *bmp6* | ZDB-GENE-050306-42 | (Thisse and Thisse, 2004) |
| *bmp7b* | ZDB-GENE-060929-328 | (Shawi and Serluca, 2008) |
| *bmper* | ZDB-GENE-030219-146 | (Rentzsch et al., 2006) |
| *cdh5* | ZDB-GENE-040816-1 | (Larson et al., 2004) |
| *chordin* | ZDB-GENE-990415-33 | (Hammerschmidt et al., 1996) |
| *dacha* | ZDB-GENE-020402-3 | (Hammond et al., 2002) |
| *dlx3b* | ZDB-GENE-980526-280 | (Ekker et al., 1992) |
| *dlx5a* | ZDB-GENE-990415-49 | (Akimenko et al., 1994) |
| *fn1a* | ZDB-GENE-000426-1 | (Trinh and Stainier, 2004) |
| *foxd3* | ZDB-GENE-980526-143 | (Odenthal and Nüsslein-Volhard, 1998) |
| *foxi1* | ZDB-GENE-030505-1 | (Solomon et al., 2003) |
| *gata2a* | ZDB-GENE-980526-260 | (Detrich et al., 1995) |
| *gbx2* | ZDB-GENE-020509-2 | (Rhinn et al., 2003) |
| *hmx2a* | ZDB-GENE-080506-2 | (Feng and Xu, 2010) |
| *hmx3a* | DB-GENE-001020-1 | (Feng and Xu, 2010) |
| *lmo4b* | ZDB-GENE-030131-3570 | (Thisse and Thisse, 2004) |
| *lmx1ba* | ZDB-GENE-050114-3 | (McMahon et al., 2009) |
| *lmx1ba* | ZDB-GENE-050114-2 | (Obholzer et al., 2012) |
| *mbp* | ZDB-GENE-030128-2 | (Brösamle and Halpern, 2002) |
| *Neurod1* | ZDB-GENE-990415-172 | (Blader et al., 1997) |
| *otx1b* | ZDB-GENE-980526-400 | (Li et al., 1994) |
| *pax2a* | ZDB-GENE-990415-8 | (Pfeffer et al., 1998) |
| *pcna* | ZDB-GENE-000210-8 | (Koudijs et al., 2005) |
| *prdm1a* | ZDB-GENE-030131-2193 | (Baxendale et al., 2004) |
| *smad6b* | ZDB-GENE-050419-198 | (Thisse and Thisse, 2004) |
| *sox10* | ZDB-GENE-011207-1 | (Dutton et al., 2001) |
| *tecta* | ZDB-GENE-110411-120 | (Stooke-Vaughan et al., 2015) |
| *ugdh* | ZDB-GENE-011022-1 | (Busch-Nentwich et al., 2004) |
| *vcanb* | ZDB-GENE-030131-2185 | (Kang et al., 2004) |
